# Long-range mRNA folding shapes expression and sequence of bacterial genes

**DOI:** 10.1101/2025.11.27.690947

**Authors:** Manraj S. Gill, Isabelle A. Kim, James R. Xue, Yashna Thappeta, James C. Taggart, Gene-Wei Li

## Abstract

Accessibility of the ribosome binding site (RBS) plays an outsized role in bacterial mRNA decay and translation. Antagonistic mRNA sequences that reduce accessibility and regulate expression have been widely documented near the RBS. To determine whether such sequences are also the primary effectors of expression when placed far from the RBS, we measured impacts of all possible 8-nucleotide substitutions (65,536 variants) at different positions in mRNA in *Bacillus subtilis*. While the vast majority of substitutions negligibly affect RNA levels, pyrimidine-rich substitutions resembling the anti-Shine-Dalgarno (aSD) sequence exhibit strong inhibitory effects. Even several hundred nucleotides downstream of the RBS, these aSD-like sequences base-pair with the RBS, promote RNA decay, and inhibit translation initiation. We find aSD-like sequences to be depleted throughout endogenous genes, likely due to selective pressure for expression. Taken together, our findings reveal widespread long-range RNA intramolecular interactions *in vivo* and uncover a key constraint on gene sequence evolution.

**HIGHLIGHTS:**

- Long-range mRNA folding tunes accessibility of the ribosome binding site (RBS)
- Short anti-RBS sequences are major mRNA repressors across the transcript body
- Anti-RBSs, even distally located, can promote RNA decay and inhibit translation
- Anti-RBS sequences are depleted throughout endogenous bacterial coding sequences

## INTRODUCTION

mRNA secondary structures are critical determinants of bacterial gene expression. Structural elements (e.g., stem loops, pseudoknots, and G-quadruplexes) can drastically alter mRNA transcription, degradation, or translation (Sharp, 2009; Mortimer et al., 2014; Del Campo et al., 2015). Many regulatory features are composed of these structural elements and include intrinsic transcription terminators, riboswitches, and endoribonucleolytic cleavage sites (Breaker, 2012; Ray-Soni et al., 2016; Altuvia et al., 2018; Taggart et al., 2025). Most of these features known to date are short (up to 200 nucleotides in length), potentially due to limitations of *in silico* structure predictions and *in vivo* structure probing (Doshi et al., 2004; Lange et al., 2012; Mustoe et al., 2018; Marinus et al., 2021; Flamm et al., 2022; Sato and Hamada, 2023; Schneider et al., 2023; Allan et al., 2024). A fundamental open question is whether long-range intramolecular interactions within an mRNA molecule are functionally consequential *in vivo*.

A major hub of regulation through RNA structure is the accessibility of the translation initiation site to ribosomes. A typical ribosome binding site (RBS) spans ∼30 nt, including a start codon and a consensus Shine-Dalgarno (SD) sequence 5′-AGGAGG-3′ (Shine and Dalgarno, 1974; Chen et al., 1994). The SD region recruits the 30S ribosomal particle by base pairing to the pyrimidine-rich anti-SD (aSD) site 5′-CCUCCU-3′ of the 16S RNA (Steitz and Jakes, 1975). Numerous *cis*-elements in the 5′ <u>u</u>ntranslated <u>r</u>egion (UTR) modulate RBS accessibility via specific RNA secondary structures that compete with ribosome binding (McCarthy and Gualerzi, 1990, de Smit and van Duin, 1990a; de Smit and van Duin, 1990b; de Smit and van Duin, 1994), such as riboswitches (Breaker, 2018) and RNA thermometers (Kortmann and Narberhaus, 2012).

The accessibility of the RBS impacts mRNA decay in addition to the rate of translation initiation. Ribosomes near the 5′ end of mRNAs can impede the activity of various RNA degradation machineries, including 5′ pyrophosphohydrolases (RppH) and 5′ end-dependent endo- and exo-ribonucleases (e.g., RNases E, Y, and J) (Agaisse and Lereclus, 1996; Sharp and Bechhofer, 2005; Richards et al., 2012; Richards and Belasco, 2019; Cetnar and Salis, 2021; Cetnar et al., 2024). Importantly, this mRNA stabilizing effect can be achieved by ribosome binding near the 5′ end, even without translation initiation (Hambraeus et al., 2002; Sharp and Becchofer, 2003).

Consistent with RBS accessibility exerting a dramatic effect on protein synthesis, the surrounding sequences (near the mRNA 5′ end) appear to be under strong selection. For native mRNAs, structural probing experiments and computational analyses demonstrated that the RBS regions are substantially less structured compared to coding sequences (Gu et al, 2010; Scharff et al., 2011; Burkhardt et al., 2017; Mustoe et al., 2018). For synthetic mRNAs, massively parallel reporter assays showed that synonymous substitutions of the first few codons can suppress protein synthesis by base pairing with the RBS (Kudla et al., 2009; Goodman et al., 2013; Bhattacharyya et al., 2018; Borujeni and Salis, 2016; Borujeni et al., 2017). These results corroborate the enrichment of rare codons in the N-termini of bacterial genes, which is computationally predicted to reduce base-pairing with the RBS (Eyre-Walker and Bulmer, 1993; Tuller et al., 2010, Bentele et al., 2013). However, extending such analyses beyond the N-terminal codons has been challenging due to the difficulty in predicting long-range RNA folding (Kelsic et al., 2016; Umu et al., 2016).

Long-range RNA folding can occur *in vivo*, but its relevance in bacterial mRNAs remains understudied. Large-scale structures are common to non-coding RNAs and viral RNAs (Hahn et al., 1987; Cruz and Westhof, 2009; Polacek et al., 2009; Nicholson and White, 2015; Palo et al., 2025). In bacteria, several reports of mRNA base pairing between the 3′ and the 5′ UTR (*hbs* in *B. subtilis* and *icaR* in *S. aureus*) suggest that distal ends of mRNA can interact and have functional consequences (Ruiz de los Mozos et al., 2013; Braun et al., 2017). Together, these lines of evidence suggest that bacterial mRNAs may also exhibit long-range RNA folding. If long-range mRNA folding indeed has major impacts on protein synthesis, selection on expression level could influence the evolution of coding sequences.

In this study, we examine the impact of RBS-complementary sequences across different regions of the mRNA. We utilize *B. subtilis* because its uncoupled transcription and translation allows us to modulate ribosome binding without affecting transcription (Johnson et al., 2020). We show that amongst all possible 8-nt sequences, those complementary to the RBS are amongst the strongest repressors of mRNA abundance — even when placed deep inside the CDS or the 3′ UTR. We further provide experimental evidence that distal anti-RBS sequences can reduce RBS accessibility, increase mRNA degradation, and inhibit translation. Lastly, we show that aSD-like sequences are depleted from both *B. subtilis* and *E. coli* coding sequences, likely due to selective pressure for expression. Taken together, our findings reveal that functional mRNA secondary structures extend beyond local interactions and impose a key constraint on gene sequence evolution.

## RESULTS

### aSD-like 8mer sequences are a major repressor of mRNA levels

To systematically examine the impact of nucleotide substitution on gene expression, we developed a <u>m</u>assively <u>p</u>arallel <u>r</u>eporter <u>a</u>ssay (MPRA) to quantify the expression level of mRNAs containing all possible 8-nt sequences (65,536 8mers) at various positions in a transcript. We engineered a reporter in *B. subtilis* based on the *aprE* transcript, which has a long half-life, making its level sensitive to variants that promote decay (Figures S1A-S1C; Hambraeus et al., 2000). Distinct MPRA libraires were built for 8mers placed at specific positions in the 5′ UTR (25 nucleotides upstream of the start codon) and coding sequence (467 nt downstream) (Figures 1A, S1D, and Methods). The RNA level for each variant was measured using deep sequencing as its frequency in the RNA pool normalized by its frequency in the DNA pool (Figures S2A-S2C). We further optimized the library preparation protocol to minimize measurement biases generated by reverse transcription and PCR (Figures S3A-S3J and Methods).

**Figure 1.**
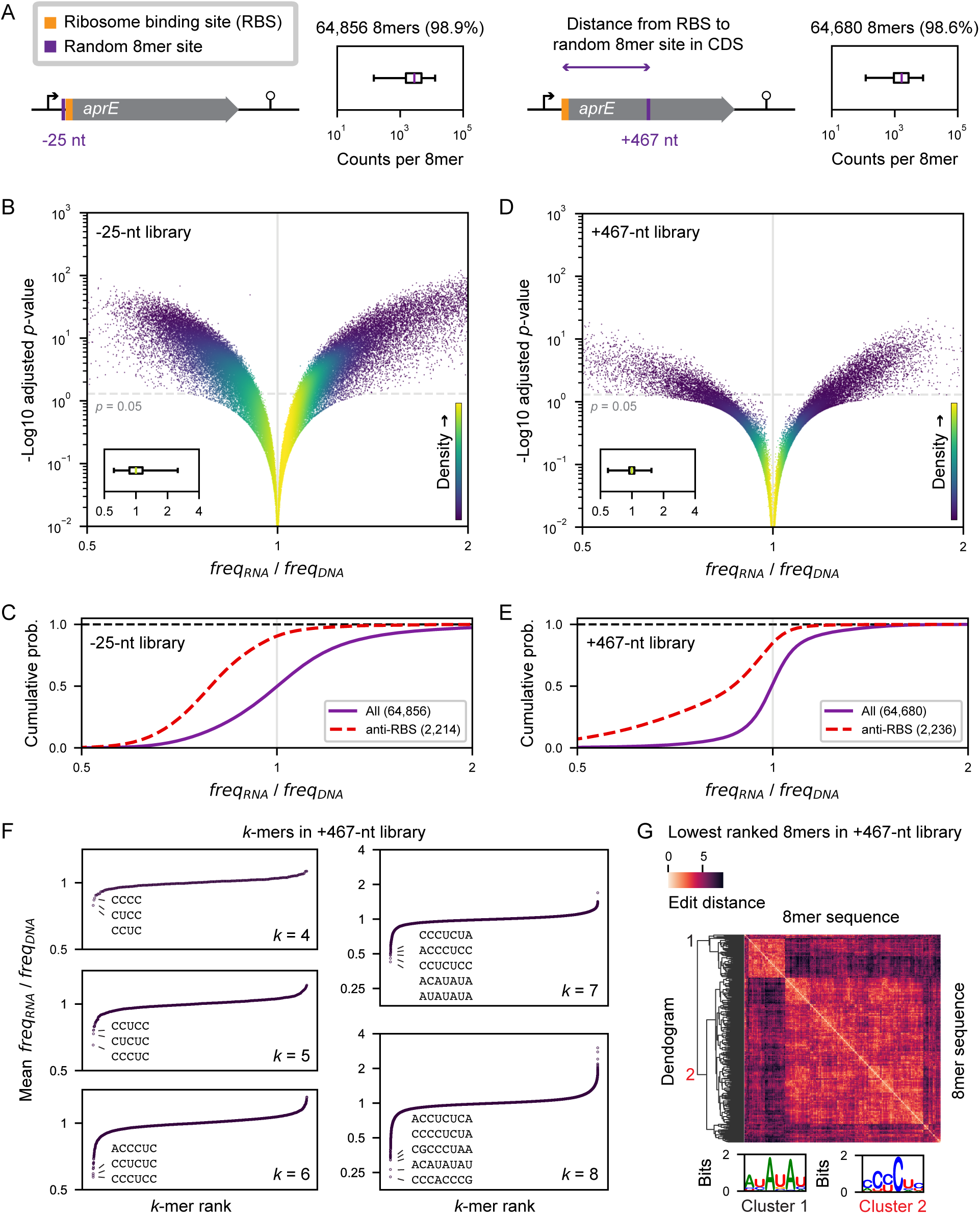
8mer sequences complementary to RBS are a major repressor of mRNA level, whether placed in 5′ UTR or deep inside the CDS. (**A**) Schematic of 5′ UTR and CDS MPRA libraries showing the −25-nt and +467-nt sites where the random 8-nt substitutions are introduced. Coding sequences drawn to scale and flanked by transcriptional start and end points. Inset: Composition of the libraries analyzed with the percent of all possible 8mers above 50 gDNA sequencing read counts denoted. Total gDNA counts of 226M were obtained for the −25-nt library and 133M for the +467-nt library. Middle line is the median, box extends from Q1 to Q3 and the whiskers extend from the 1st to the 99th percentiles. (**B**) Steady-state level of mRNA variants with 8mers in the 5′ UTR shown as the freq_RNA_/freq_DNA_ normalized around the median in the library vs. their corresponding Benjamini-Hochberg adjusted *p*-value. Colors correspond to kernel density estimation of the number of variants per pixel. Inset: Distribution of freq_RNA_/freq_DNA_ observed, middle line is the median, box extends from Q1 to Q3 and the whiskers extend from the 1st to the 99th percentiles. (**C**) Cumulative distribution of the relative mRNA level of 5′ UTR 8mers with sequence complementary to the RBS (anti-RBS; free energy of binding to the RBS less than −8 kcal/mol), compared to the distribution of all variants shown in **B**. (**D-E**) Same as in **B-C** for 8mers at the +467-nt site in the CDS. (**F**) *k-*mer analysis is shown as the mean level of *k* = 4, 5, 6, 7, and 8 from the +467-nt library, after accounting for *k*-1 nucleotides of the flanking sequence on either side and ranking in ascending order of mean effect. Lowest ranked *k-*mers from each set are highlighted. (**G**) Clustering based on the Levenshtein edit-distance between each 8mer of the 500 lowest ranked 8mers from the +467-nt library. Motif enrichment analysis on sequences from each cluster is shown as the inset. 104 8mers in cluster 1 and 396 in cluster 2.

We expected that in the 5′ UTR, 8mers that affect RBS accessibility would have the strongest effects on expression levels. Among the 64,856 quantifiable variants in our 5′ UTR −25-nt library (98.9% of the possible sequences; Figure 1A), ∼75% have negligible effects on the steady-state mRNA levels (0.75 < freq_RNA_/freq_DNA_ < 1.25, relative to the median 5′ UTR 8mer; Figure 1B). As expected, 8mers complementary to the RBS lower mRNA levels (anti-RBS, Figure 1C). Conversely, the 8mers that significantly increase mRNA levels are predicted to increase the accessibility of the RBS (Figure S4A). These results confirm that variants in the vicinity of RBS have strong effects on mRNA levels and validate our approach to detect systematic effects.

Surprisingly, when we placed the 8mer library in the CDS (+467-nt library, Figure 1A), the 8mers complementary to the RBS also significantly decreased RNA levels. Among the 64,680 variants (98.6% of the possible sequences), ∼90% have negligible effects (0.75 < freq_RNA_/freq_DNA_ < 1.25, relative to the median CDS 8mer; Figure 1D). However, anti-RBS 8mers strongly lower mRNA levels (Figure 1E). To determine sequence features underlying the CDS MPRA trends, we analyzed *k-*mer subsequences that included the nucleotides flanking the 8mer site and ranked each *k*-mer by its mean effect. Pyrimidine-rich *k*-mers complementary to the consensus SD have particularly strong effects in reducing RNA levels, and this effect strengthens with increasing length (Figure 1F). Furthermore, when we clustered the lowest ranked 8mers by their nucleotide similarity (Levenshtein edit distance), the aSD-like pyrimidine-rich sequences represent the largest cluster (Figure 1G). An additional smaller cluster shares a 5′-AUAUAU-3′ motif and may present an endonucleolytic cleavage site or another repressive element of gene expression. Taken together, we discovered that aSD-like sequences are major repressors of mRNA levels even deep in the CDS.

### aSD-like 8mers in CDS promote mRNA degradation via long-range folding

The internal aSD-like 8mers could decrease mRNA levels either by abolishing transcription or promoting mRNA degradation. To directly measure mRNA stability, we induced reporter expression and measured the time to reach half its steady-state level. This half-life upon induction is inversely proportional to the degradation rate and is independent of transcription rates (Methods; Alon, 2007). We tested one of the lower ranked +467-nt site 8mers (denoted as aSD*: 5′- CCCUCUCC-3′; freq_RNA_/freq_DNA_ = 0.31) that is perfectly complementary to the SD region of the *aprE* reporter (5′-GGAGAGGG-3′). We found that the aSD* 8mer substantially shortened the half-life (1.7 ± 0.2 minutes compared to 11.2 ± 0.8 minutes for the WT 8mer, Figure 2A), matching the reduction in steady-state mRNA level. These results established that the aSD* 8mer decreases mRNA level by increasing its degradation rate.

**Figure 2.**
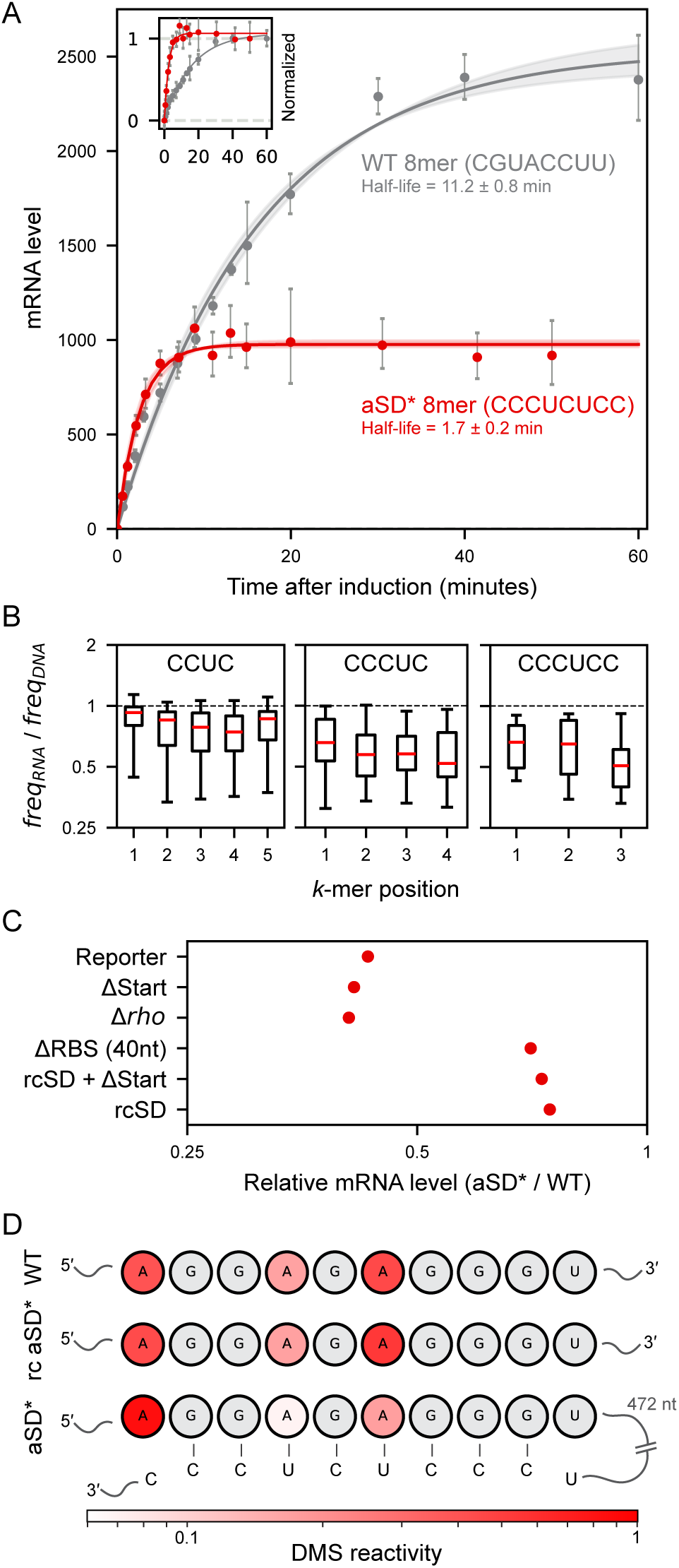
RBS-complementary sequences in CDS engage in long-range mRNA folding, which decreases mRNA level by promoting its degradation. (**A**) Half-life measurements from induction curves of reporters with the WT 8mer sequence and the aSD* 8mer sequence in the +467-nt CDS site measured by RT-qPCR. mRNA level shown is relative to 0 min pre-induction. Inset: induction curves with mRNA level normalized to 60 min. Error bars represent propagated standard error from 3 technical replicates and shaded region represents the 95% confidence interval of the fit. (**B**) Reading-frame analysis of lowest ranked *k*-mers from **1F** across their possible registers in the +467-nt site. *k*-mer position 2 is the first nucleotide of the translational reading frame for all. (**C**) Steady-state mRNA levels from RT-qPCR measurements measuring the effect of the aSD* 8mer relative to the WT sequence at the +467-nt site, across mutations elsewhere in the reporter or genome. Each point represents the mean of 3 biological replicates. Δ denotes deletion of the start codon (GUG), the *rho* gene, or the full RBS (spanning from −17 to +23 relative to the start codon). The SD region (5′- AGGAGAGGG-3′) is mutated to its reverse complement (5′- CCCUCUCCU-3′) in reverse complement SD (rcSD). (**D**) Normalized DMS-MaPseq mutation rate (DMS reactivity) of As in the SD region with the WT, reverse complement aSD* (rc aSD*), and aSD* at the +467-nt CDS site. Higher values correspond to greater base accessibility. Grey bases (Gs and Us) are not susceptible to DMS mutation. The sequence of the aSD* 8mer (5′-CCCUCUCC-3′), only present in the aSD* variant, is highlighted at the bottom to show its complementarity with the SD region.

Multiple mechanisms can promote mRNA degradation, so we next ruled out several potential contributions from changes in translational and transcriptional elongation. First, the effects of the strongest *k*-mers are similar across different reading frames, suggesting that degradation is not triggered by changes in translation elongation (Figure 2B). Second, although the pyrimidine-rich aSD-like sequences can resemble groups of proline codons (CCN) that could stall translation and subsequently destabilize mRNA, the effects of CCN and CCNCCN are also independent of the reading frame (Figures S4B-C). Third, the effect is independent of any ribosome traffic across the CDS, as mutating the start codon of the reporter to a stop codon does not affect how aSD* increases mRNA degradation (ΔStart, Figure 2C). Finally, although the pyrimidine-rich aSD-like sequences also resemble the transcription termination signal recognized by Rho (Ciampi, 2006; Dierksheide et al., 2025), we found that the destabilizing effect is independent of the presence of Rho (Δ*rho*, Fig. 2C).

Next, we showed that aSD-like sequences promote mRNA degradation by engaging in long-range mRNA folding specifically with the SD sequence in the 5′ UTR. To test this hypothesis, we measured the fold-change between the WT 8mer and the aSD* 8mer in various RBS mutants of the reporter transcript. We observed that the degradation effect of the aSD* 8mer is diminished in a deletion of the RBS (ΔRBS, Figure 2C). Moreover, this effect is specifically confined to the SD region of the RBS, as replacing the SD with its reverse complement (rcSD), is sufficient to diminish the degradation effect of adding the aSD* to the CDS (Figure 2C). These results suggest that the faster decay rate of the transcript caused by the aSD* 8mer in the +457-nt site is specifically dependent on its base-pairing interaction with the SD.

To experimentally determine SD accessibility *in vivo*, we used the RNA structure probing method targeted <u>d</u>i<u>m</u>ethyl <u>s</u>ulfate-<u>m</u>ut<u>a</u>tional <u>p</u>rofiling and <u>seq</u>uencing (DMS-MaPseq). In DMS-MaPseq, the A and C bases of RNA molecules exhibit a higher mutation rate when they are unpaired. With aSD* (5′-CCCUCUCC-3′) at +467, both As in the SD region (5′-GGAGAGGG-3′) exhibit a lower mutation rate than reporters with the wildtype sequence or the reverse complement of aSD*. These results indicate that the distal aSD* shielded the SD region from DMS modification by base pairing (Figure 2D). Computational folding of this mRNA, constrained by DMS reactivity surrounding the SD region (a 157-nt window), further supports that the SD sequence is base paired with the aSD* at +467 via long-range RNA folding (Figure S5).

Taken together, the results from reporter mutants and *in vivo* structure probing suggest that the aSD-like 8mers promote mRNA degradation by engaging in long-range mRNA folding with the SD, leading to decreased accessibility of the ribosome to the RBS, a lowered 5′ UTR occupancy of the ribosome, and thereby less protection from the ribosome against the activity of 5′-end dependent mRNA degradation machinery.

### Long-range mRNA folding by aSD-like sequences blocks translation

Given that internal aSD-like sequences destabilize mRNA by altering ribosome binding, we hypothesized that such sequences can also directly inhibit translation from a distance. To measure this effect systematically, we designed a series of translational reporters with an aSD-like 12mer placed at varying positions along the transcript (Figure 3A). The aSD-like 12mer is complementary to the reporter’s SD sequence and following 6 nucleotides. Rather than using the *aprE* reporter that produces secreted proteins, we fused a cytosolic protein (Ctc) with a fluorescent protein (mNeonGreen) and used intracellular fluorescence to measure the effect of internal aSD-like sequences on protein synthesis (Figure 3A). We also included one of *ctc*’s native 5′ UTRs that is less structured and highly translated (McCormick et al., 2021), enabling us to probe the impact of RBS base-pairing mediated by downstream sequences. Translation efficiency for each variant was calculated by dividing the fluorescence signal of each strain by its mRNA level (Figure 3A).

**Figure 3.**
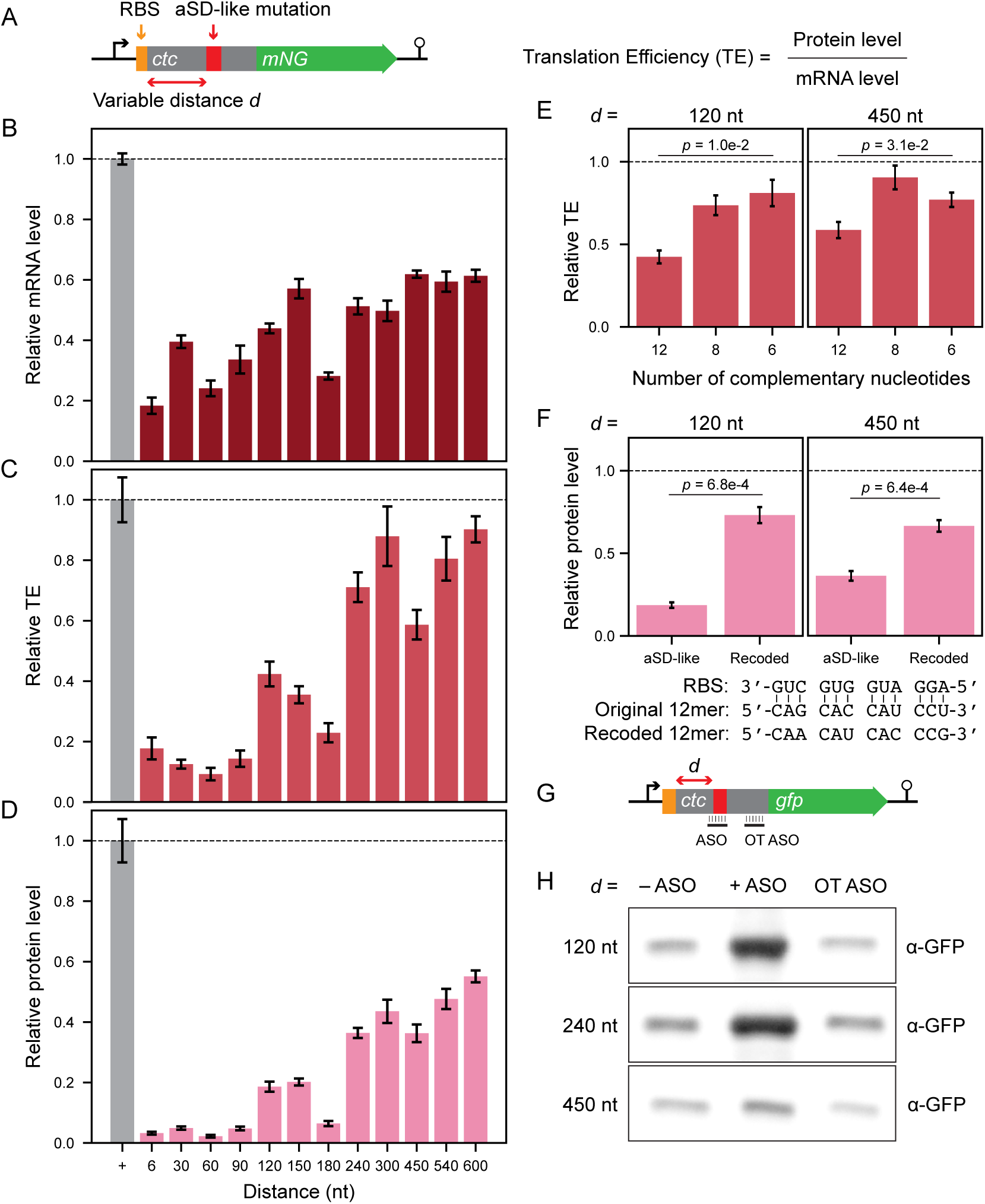
Long-range mRNA folding can block ribosome binding. (**A**) Schematic of *ctc-mNG* translation efficiency reporter (not to scale). Highlighted regions include a constant RBS in the 5′ UTR and an aSD-like mutation at different distances (*d*) from the RBS. The aSD-like mutation is complementary to the SD sequence and downstream nucleotides. The positive control is the native *ctc* sequence and does not contain an aSD-like mutation. Translation efficiency (TE) of each reporter variant is calculated by dividing protein level by mRNA level. Relative protein level is measured by mNG fluorescence normalized to OD and mRNA is measured using RT-qPCR. (**B**) Relative mRNA level of different *ctc-mNG* reporters with aSD-like 12mers normalized to the positive control. Error bars represent the standard error of the mean of three biological replicates with four technical replicates each for the positive control, and four technical replicates for all other variants. (**C**) Relative translation efficiency (TE) of different *ctc-mNG* reporters with aSD-like 12mers normalized to the positive control. Error bars represent the standard error of the mean of three biological replicates with four technical replicates each for the positive control, and four technical replicates for all other variants. (**D**) Relative protein level of different *ctc-mNG* reporters with aSD-like 12mers normalized to the positive control. Error bars represent the standard error of the mean of three biological replicates with four technical replicates each for the positive control, and four technical replicates for all other variants. (**E**) Relative TE for reporters with 12, 8, and 6 nucleotides of complementarity between the RBS and aSD-like sequence for variants with 120 and 450 nucleotides of distance between the start codon and anti-RBS sequence. *p-*values were obtained from a two-sided Student’s *t*-test. (**F**) Relative protein levels for reporters whose aSD-like 12mers have undergone synonymous mutations to weaken base pairing interactions with the RBS, but maintain the amino acid identity, for strains with 120 and 450 nucleotides of distance between the start codon and anti-RBS sequence. *p-*values were obtained from a two-sided Student’s *t*-test. The recoded sequence decreases the complementarity to the RBS from 12 base pairs to 9. (**G**) Schematic of *in vitro* translated reporter (not to scale). The native *ctc* 5′ UTR and CDS are fused to *gfp* with a linker. aSD-like 12mers are substituted in at a variable distance *d* from the *ctc* start codon and antisense oligonucleotides (ASOs) bind either over the aSD-like 12mer or at an off-target site on *ctc.* ASOs ranged from 18-26 nt to achieve a T_m_ of 72-73 °C in reaction conditions. (**H**) Anti-GFP Western blot showing the protein level of *in vitro* translated reporter mRNA without ASO, with 100-fold molar excess of ASO, and with 100-fold molar excess of off-target ASO (OT ASO) for reporters with the aSD-like 12mer located 120, 240, and 450 nt from the *ctc* start codon.

This series of reporters confirmed that aSD-like sequences effectively lower both mRNA level and translation when placed inside the CDS (Figures 3B, C). Within the first 90 nucleotides of *ctc*, 12mers that are complementary to the SD caused a nearly 10-fold decrease in translation efficiency and a 3-fold reduction in mRNA level, leading to a 30-fold reduction in protein level (Figure 3D). The inhibitory effect decreased as the distance from the start codon increased, as expected from the increased entropic cost for closing the extended RNA loop. Nevertheless, even at >500 nt downstream of the start codon, the aSD-like sequence still markedly reduced protein level by 44% (Figure 3D).

Several lines of evidence further support that the attenuated fluorescence is due to interaction between the SD and internal aSD-like sequences. First, decreasing the number of complementary nucleotides in the aSD-like sequence generally decreases the inhibitory effect of the sequence on translation at different locations along the CDS (Figure 3E). Second, synonymous mutations in the aSD-like 12mers that reduce the number of complementary nucleotides with the RBS from 12 to 9 can increase protein level in multiple locations along the transcript (Figure 3F), indicating that the attenuated fluorescence is not due to changes in the protein sequence. Third, we utilized *in vitro* translation to show that antisense oligonucleotides that specifically bind the aSD-like 12mer can rescue translation (Figures 3 G, H, and S6). Finally, DMS-MaPseq for variants with the aSD-like 12mer showed reduced accessibility of the RBS, supporting the model of base pairing between these two regions (Figure S7). Together, these results suggest that long-range intramolecular interactions can finely tune translation efficiency of native mRNAs.

### aSD-like sequences are depleted from bacterial coding sequences

The tendency of mRNAs to engage in long-range interactions, together with the experimental observation that the RBS is generally unfolded for endogenous genes, suggests that aSD-like sequences have been selected against in bacterial CDSs. To quantify the depletion of these sequences, we measured their frequency of occurrences in endogenous CDSs and compared it with a control set of randomly sampled 8mers. We observed that aSD-like 8mers are strongly depleted across *B. subtilis* genes (Figure 4A). Furthermore, although the depletion is strongest near the start codon (Figure 4B, inset), the effect persists for several hundred nucleotides (Figure 4B). We observed a similar trend for genes in monocistronic mRNAs, suggesting that the effect is not inflated by the presence of downstream RBS (Figures S8A-B; Herzel et al., 2022). Similarly for *E. coli*, which is evolutionarily separated from *B. subtilis* by >2 billion years and has a very different GC content, we observed a significant depletion in the frequency aSD-like 8mers (Figure 4C).

**Figure 4.**
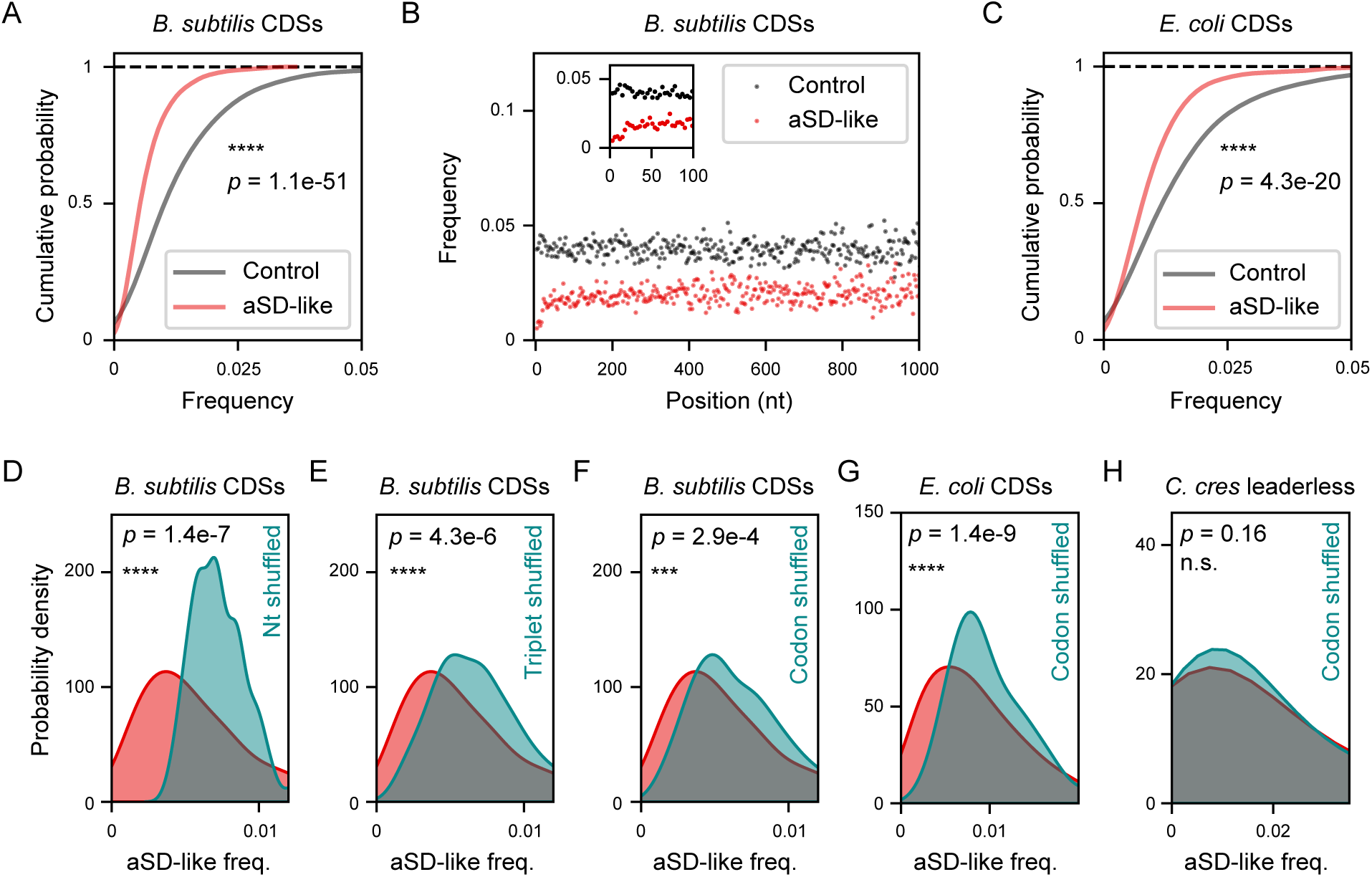
RBS-complementary sequences are depleted from bacterial coding sequences. (**A**) Frequency of aSD-like 8mers (set size = 882; free energy of binding to the consensus SD less than −8 kcal/mol) in endogenous *B. subtilis* CDSs (n = 4,218) compared to frequency of an equally sized set of control 8mers randomly sampled. *p-*value was computed from a two-sided Mann-Whitney *U* test. (**B**) Positional frequency over 1000 nt, 3-nt bins, of the frequency of aSD-like 8mers in endogenous *B. subtilis* CDSs compared to the control from **A**. Inset: 100-nt window. (**C**) Same as **A** for endogenous *E. coli* CDSs (n = 4,146). (**D-F**) Frequency of aSD-like 8mers in *B. subtilis* CDSs compared to their frequency in nucleotide shuffled CDSs (**D**), triplet shuffled CDSs (**E**), and codon shuffled CDSs (**F**). *p*-values were computed from paired two-sided Student’s *t*-tests. Mean frequencies from 20 independent shuffling instances are reported. (**G-H**) Same as **F** for *E. coli* CDSs (**G**) and leaderless CDS of *C. crescentus* (n = 374) (**H**).

The depletion of aSD-like 8mers from CDSs is not driven by biases in nucleotide frequency or codon choice. Although *B. subtilis* genes are purine-rich, the pyrimidine-rich aSD-like 8mers are more depleted in endogenous CDSs than in randomly shuffled nucleotide sequences (Figure 4D). We also controlled for overall codon composition by either randomly shuffling triplet nucleotides from endogenous genes or synonymously recoding native proteins while maintaining peptide sequences (codon shuffling; Katz and Burge, 2003). In both cases, we still observed a depletion of aSD-like 8mers in native CDSs relative to their shuffled counterparts (Figures 4E-G). These results further support our hypothesis that the depletion of aSD-like sequences throughout CDSs is driven by selective pressure to avoid less productive transcripts.

Unlike *E. coli* and *B. subtilis*, some bacterial species contain many mRNAs that do not have SD sequences or 5′ UTRs altogether, and we expect that these “leaderless mRNAs” would not be depleted of aSD-like sequences. Indeed, among the large number of leaderless mRNAs in *C. crescentus* (Schrader et al., 2014; Bharmal et al., 2021), aSD-like sequences appear at a similar frequency in their CDSs relative to codon shuffled ones (Figure 4H). Taken together, these results suggest that the presence of SD sequence in the 5′ UTR has constrained the sequence evolution of downstream genes.

### Impact of aSD-like sequences can be modulated by neighboring sequence

Many additional constraints can shape gene sequences, and we expect that their impacts are interdependent on each other. To test whether the effects of aSD-like sequences on expression can be modulated by surrounding sequence features, we created MPRA libraries that have random 8mers placed at two distinct locations in the 3′ UTR of an identical reporter. In one library (*gfp* +831-nt library with 99.9% of the possible sequences; Figure 5A), we found negative effects of anti-RBS 8mers on mRNA level (Figure 5B-C), broadly recapitulating our observations for the 5′ UTR and CDS libraries. However, at the other nearby site (*gfp* +800-nt library with 100% of the possible sequences; Figure 5D), anti-RBS 8mers have highly reduced impacts on mRNA levels, as with most other 8mers (99.9% of the 8mers with 0.75 < freq_RNA_/freq_DNA_ < 1.25 relative to the median, Figures 5E-F). In the latter site, the 8mers are placed in between two segments that could base-pair with each other, and the resulting secondary structure may shield the effects of anti-RBS 8mers. Similarly, anti-RBS sequences show no effects when flanked by two hairpin structures in the *aprE* 3′ UTR (*aprE* +997-nt library; Figures 5G-I). Therefore, while anti-RBS sequences consistently rank among the strongest repressors of mRNA level, their effect can be further modulated by surrounding sequence features, making mRNA sequence evolution a multifaceted optimization problem.

**Figure 5.**
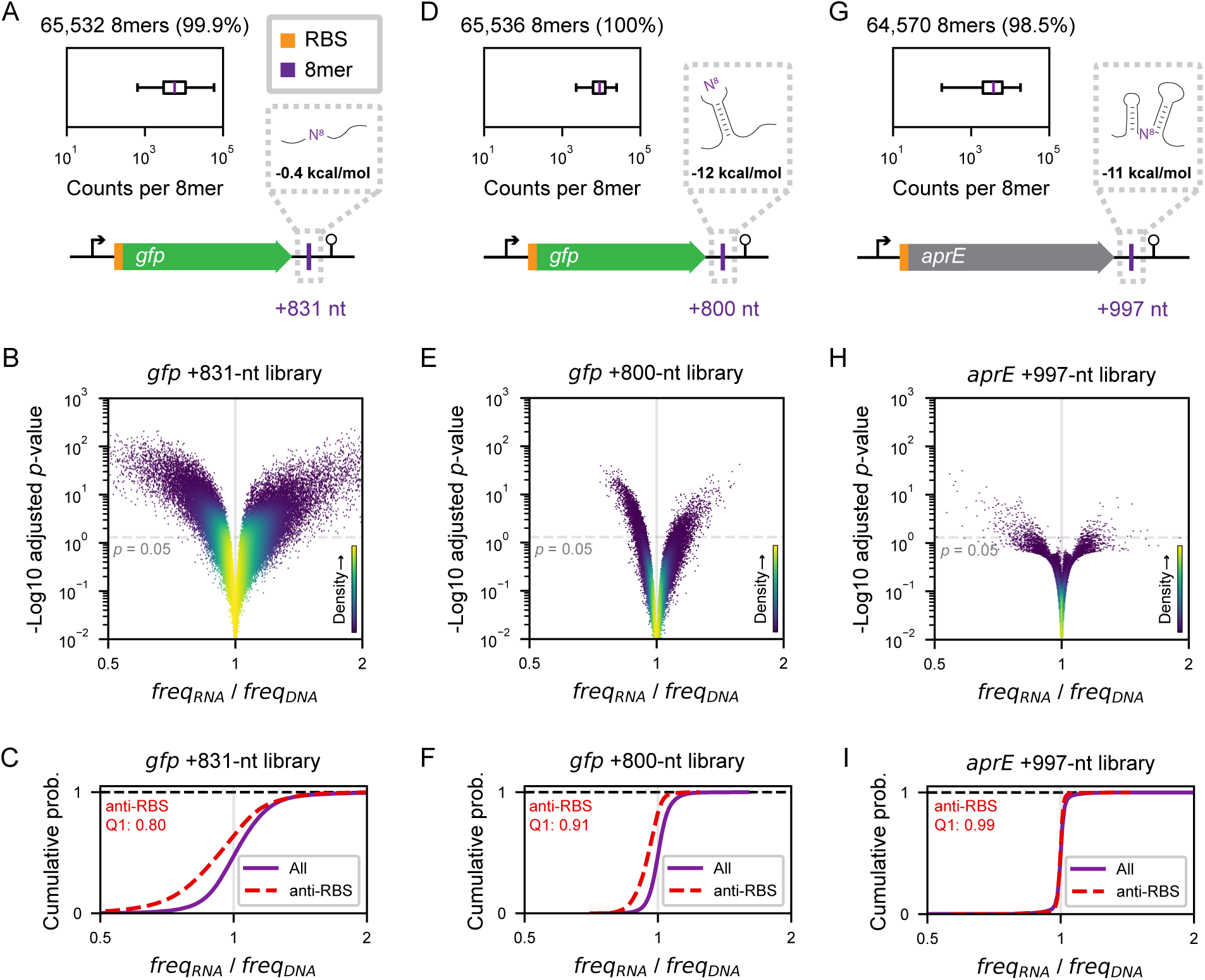
Impact of RBS-complementary sequences can be modulated by neighboring sequence. (**A**) Schematic of *gfp* 3′ UTR MPRA library showing the +831-nt site where the random 8-nt substitutions are introduced. Coding sequence drawn to scale and flanked by transcriptional start and end points. Inset: Composition of the library analyzed with the percent of all possible 8mers above 50 gDNA sequencing read counts denoted. Total gDNA counts of 645M were obtained for the +831-nt library. Middle line is the median, box extends from Q1 to Q3 and the whiskers extend from the 1st to the 99th percentiles. The centroid <u>m</u>inimum free <u>e</u>nergy (MFE) from ViennaRNA’s RNAfold of the 20-nt flanking region is also highlighted. (**B**) Steady-state level of mRNA variants with 8mers in the +831-nt site shown as the freq_RNA_/freq_DNA_ normalized around the median in the library vs. their corresponding Benjamini-Hochberg adjusted *p*-value. Colors correspond to kernel density estimation of the number of variants per pixel. (**C**) Cumulative distribution of the relative mRNA level of +831-nt site 8mers with sequence complementarity to the RBS (anti-RBS; free energy of binding to the RBS less than −8 kcal/mol), compared to the distribution of all variants shown in **B**. The 25th percentile (Q1) freq_RNA_/freq_DNA_ of the anti-RBS distribution is reported. (**D-I**) Same as **A-C** for the *gfp* +800-nt site (**D-F**) and the *aprE* +997-nt site (**G-I**).

## DISCUSSION

Our findings that long-range mRNA folding can promote decay and inhibit translation reveal a major consideration for the evolution of bacterial gene sequences (Figures 6A-D). While it is established that 5’ proximal regions can modulate RBS accessibility, our approaches circumvent the limitations of computational RNA folding, demonstrating the importance of the entire mRNA sequence. Indeed, the depletion of short aSD-like sequences from CDSs suggests that there has been selection against such sequences in bacterial genes. Therefore, a coding sequence is not just a compilation of codons, but also the driver for mRNA-wide folding that determines its own expression.

**Figure 6.**
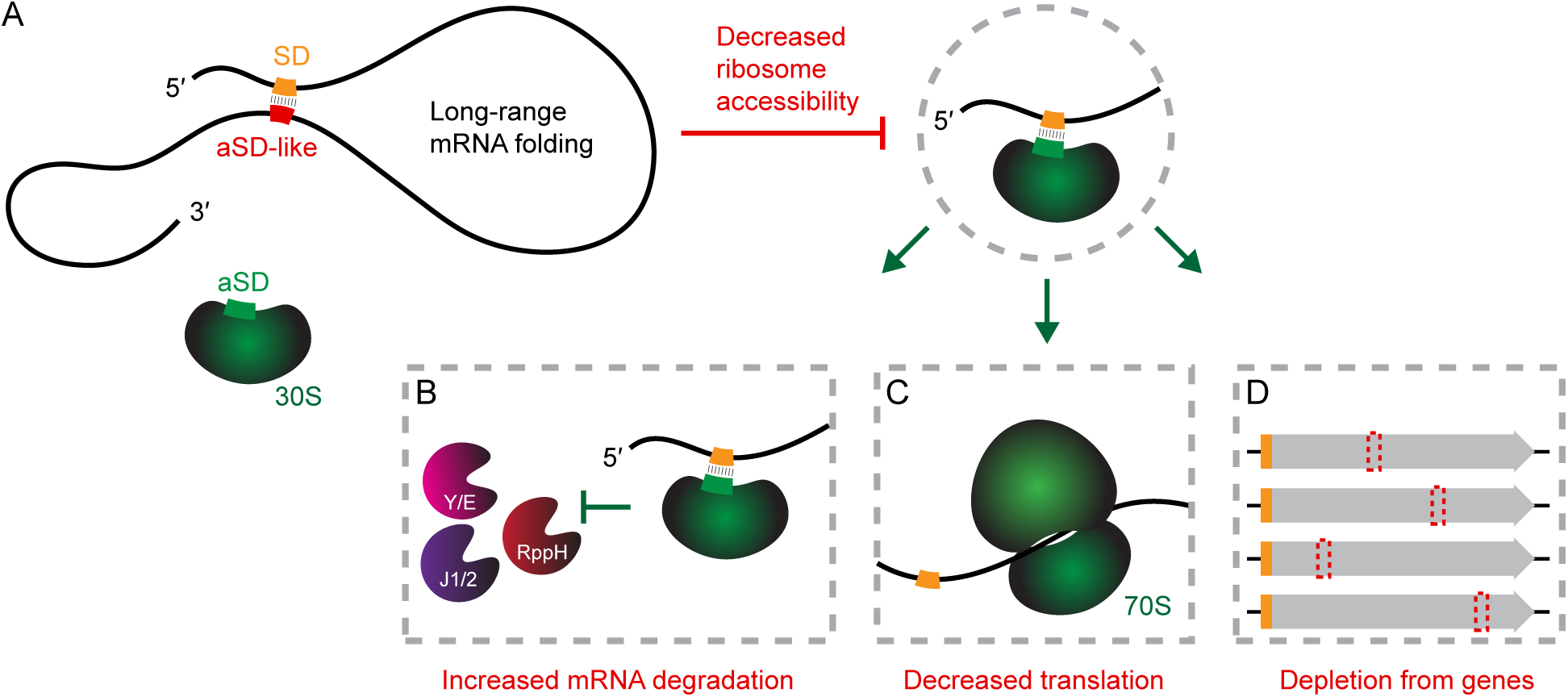
Model of long-range mRNA folding by RBS-complementary sequences shaping expression and sequence of bacterial genes. (**A**) Distal aSD-like sequences in mRNA can engage in long-range folding to decrease the accessibility of the 30S ribosomal subunit to the SD in the mRNA’s RBS. (**B**) 30S occupancy at the SD inhibits the activity of 5′-end dependent RNA degradation machinery (such as the pyrophosphohydrolase RppH, 5′-3′ exoribonucleases RNases J1/J2, and endoribonucleases RNase Y/E). Long-range folding with the SD excludes the 30S and thereby leads to increased mRNA degradation. (**C**) 30S occupancy at the SD promotes translation initiation and the subsequent formation of the 70S translation elongation complex. Long-range folding with the SD thereby leads to decreased translation. (**D**) Maintaining 30S occupancy at the SD is under selective pressure for its impacts on gene expression. Potential to interact with the SD thereby leads to depletion of aSD-like sequences throughout bacterial coding sequences.

The ability of distal mRNA regions to modulate expression may constitute a previously overlooked class of regulatory mechanisms in bacteria. Compared to compact regulatory elements that are easier to identify, few long-range interactions have been reported to date (Ruiz de los Mozos et al., 2013; Braun et al., 2017; Taggart et al., 2025). Our results suggest that the functional RNA interactome may include abundant self-interactions at a distance. Because mRNA folding kinetics are closely tied to ribosome traffic (Takyar et al., 2005), we anticipate that long-range self-interactions can be utilized to sense changes in translation elongation or initiation. Furthermore, in polycistronic mRNAs, long-range interactions may help cross-regulate or insulate neighboring genes (Burkhardt et al., 2017). Therefore, the regulatory potential for post-transcriptional processes can be embedded across the entire mRNA, extending far beyond the vicinity of the ribosome binding site.

The impact of long-range mRNA folding adds to a growing list of considerations that shape gene sequences. Sequences that recruit ribonucleases (Durand et al., 2012; Bandyra and Luisi, 2018; Taggart et al., 2025), stall transcription (Yakhnin et al., 2020; Dierksheide et al., 2025), stall translation (Li et al., 2012; Ito and Chiba, 2013), and are decoded by rare tRNAs (Plotkin and Kudla, 2011) can also impact gene expression. Although long-range interactions with the RBS were found to play an outsized role, we expect the magnitude of effects to be entrenched by these other considerations. Indeed, between and even within different reporters (*aprE*, *ctc-mNG*, and *gfp*), we observed that aSD-like sequences have unequal effects on mRNA levels (Figures 3B and 5). These results highlight the need for a holistic model to predict the impact of mutations on expression.

Our results further suggest that such holistic sequence-to-expression models must overcome the current barrier in RNA structure prediction. In addition to the combinatorial complexity involved in long-range folding, RNAs *in vivo* are also influenced by the kinetics of transcription and translation, as well as a plethora of other interaction partners. Although advances in deep-learning models have proven useful to predict RNA structure and expression levels from gene sequence (Singh et al., 2019; Kappel et al., 2020; Singh et al, 2021; Fu et al., 2022; Schneider et al., 2023; Sato and Hamada, 2023; Allan et al., 2024; Wang et al., 2025), the power of these models is still limited by a dearth of high-quality training datasets measured *in vivo*. Future application of MPRAs for quantifying the impact of targeted sequence variations on mRNA stability and translation will help refine and inform deep learning models and define the RNA “structurome” (Spitale et al., 2023).

In eukaryotes and viruses, long-range messenger RNA structure has been recognized as an important consideration for gene expression, through its action in splicing, 5′–3′ interaction during translation initiation, and RNA stability (Wells et al., 1998; Yang et al., 2015; Yue et al., 2016; Xiang and Bartel, 2021; Kalmykova et al., 2021; Margasyuk et al., 2023). Our study indicates that the regulatory functions of long-range mRNA structures may be also widespread in bacteria and profoundly impact coding sequence evolution.

## RESOURCE AVAILABILITY

### Lead contact

Requests for additional information, resources, and data should be directed to and will be fulfilled by the lead contact, Gene-Wei Li.

### Materials availability

All reagents generated are listed in the key resources table on Mendeley Data and are available by request from the lead contact.

### Data and code availability

- The data for this study have been deposited in the European Nucleotide Archive (ENA) at EMBL-EBI and the accession number will be publicly available upon publication.
- All original code and supplementary tables deposited on Mendeley Data and will be publicly available upon publication.
- Any additional information required to reanalyze the data reported in this paper is available from the lead contact upon request.

## ACKNOWLEDGEMENTS

The authors thank the MIT BioMicro Center, especially Stuart Levine, Christopher Hallee, and Noelani Kamelamela for advice and high-throughput sequencing, the members of the Li lab for valuable discussion and feedback, Christopher Burge, David Bartel, Dylan McCormick, Adam Sychla, Tammy Lan, Silvi Rouskin, Julia Dierksheide, Wen Du, and Armaan Grewal. G.-W.L. is an Investigator at the Howard Hughes Medical Institute (HHMI). This work was supported by: William Asbjornsen Albert Memorial Fellowship (to M.S.G.), MathWorks Fellowship (to M.S.G.), MIT School of Science Service Fellowship (to M.S.G.), NIH Training Grant 5T32GM007287 (to M.S.G., I.A.K., and J.C.T.), Helen Hay Whitney Foundation Fellowship (to J.R.X.), and NIH R35GM124732 (to G.-W.L.).

## AUTHOR CONTRIBUTIONS

Investigation and formal analysis: M.S.G. (*aprE*, aSD-like depletion), I.A.K. (*ctc-mNG*), J.R.X. (*gfp*), Y.T. (DMS); Methodology (MPRA): M.S.G.; Resources (Strains): J.C.T.; Visualization: M.S.G., I.A.K., Y.T.; Writing - review and editing: M.S.G., I.A.K., Y.T., J.R.X., J.C.T., G.-W.L.; Writing - original draft: M.S.G., I.A.K., G.-W.L.; Supervision: G.-W.L.

## METHODS

### Plasmid construction

Plasmids for all reporters (Supplementary Table S2) were constructed through restriction cloning or Gibson assembly using the oligos listed in Supplementary Table S3. We developed *aprE* to create an inducible reporter by modifying the promoter and the 5′ end of the native *aprE* transcript to be regulated by a strong inducible promoter under the control of TetR binding (Peters et al., 2019). Addition of anhydrotetracycline (aTc) inactivates TetR binding on the promoter and induces expression. Further, the 3′ UTR was extended to create a platform for the +997-nt site by introducing a premature stop codon. Introduction of a premature stop does not lower mRNA levels, consistent with a general lack of nonsense-mediated polarity in *B. subtilis* (Johnson et al., 2020) and with an increase in mRNA levels observed from in-frame stop codons at the +467-nt site (Figures S9A-C). Additionally, an NheI (NEB) restriction site was introduced upstream of the +997-nt site and is present in all reporter variants. For cloning the 5′ UTR library, an AscI (NEB) restriction site was introduced upstream of the −25-nt site and is only present in that library. For the +467-nt library an HindIII (NEB) restriction site in the native sequence was leveraged for cloning. To construct the *aprE* MPRA plasmid libraries, empty vector variants of the reporters were purified (ZymoPURE II Plasmid Midiprep Kit) and 15 μg were digested at 37 °C with corresponding restriction enzymes (NEB). The linearized empty vectors were purified (Zymo Clean & Concentrator-5 Kit) and 1.5 μg was ligated with 750 ng of digested inserts (amplified with oligos containing the random 8mer; Figure S1D) at a 1:10 ratio using 15 μl NEB QuickLigase for 2.5 hours at 25 °C. The ligation reactions were purified (Zymo Clean & Concentrator-5 Kit) and transformed into electrocompetent *E. coli* cells (NEB 10-beta) using 1 μl per 25 μl cells across 10-15 cuvettes (Eppendorf Eporator). All cultures were combined into one flask for recovery at 37 °C for 60 minutes prior to selection in 100 mL LB with the corresponding antibiotic overnight. Plasmids were purified (ZymoPURE II Plasmid Midiprep Kit) from the resulting cell pellet and transformed into *B. subtilis*. Additional information and library-specific details are available in the extended methods sections.

To construct the strains for the fluorescent reporter assay, we utilized the plasmid pDM001 from McCormick et al., 2021 which contains the *B. subtilis* open reading frame *ctc* with a carboxy-terminal linker fused to mNeonGreen. The original plasmid’s *levB* homology regions were swapped out for *amyE* homology regions using HiFi assembly (NEB) to create pIK51. For strains with the *ctc* aSD-like 12mer in various locations in the transcript body, gene blocks containing the 12mer and flanking regions were cloned into the pIK51 backbone (Twist Biosciences). The assembled plasmids were transformed into *Mix and Go! E. coli* DH5 Alpha Competent Cells (Zymo) and isolated via QIAprep Spin Miniprep Kit (Qiagen).

### Strain construction

For all experiments, derivatives of *B. subtilis* W168 were used (Supplementary Table S1). For integration of the purified *aprE* MPRA plasmids, 30 μg of each of the plasmid libraries were linearized with ScaI-HF (NEB) at 37 °C for 75 minutes. A colony from a Δ*aprE* strain with P_tet_-*comK* at *amyE* was picked into 30 ml LB and allowed to grow to an OD600 of 1.0 at 37 °C with vigorous shaking. At this point, aTc was added (final concentration 20 ng/ml) to induce expression of ComK. Cells were shaken at 37 °C for an additional 2 hours, after which all 1.5 ml containing the 30 μg linearized library was added to 15 ml of competent cells. The cells were cultured with DNA for 1.5 hours (up to 8 hours for the *gfp* MPRA library construction) at 37 °C and then resuspended in a total volume of 120 ml LB with the corresponding antibiotic for selection overnight prior to preparation of glycerol stocks of the MPRA libraries. Other plasmid variants of *aprE*, *ctc-mNG*, and *gfp* reporters were similarly integrated into the *amyE* locus using either a standard protocol relying on natural competence of *B. subtilis* or using derivatives of the aTc inducible ComK overexpression strain described above. Transformants for each such case were routinely verified via colony PCR and Sanger sequencing (Quintara Biosciences). Additional information and library-specific details are available in the extended methods sections.

For strains containing the *ctc* reporter that were used in DMS MaPseq, the native copy of *ctc* was markerlessly removed using the CRISPR protocol described in Burby and Simmons 2017. *ctc* reporters were then transformed into the Δ*ctc* background strain using aTc inducible ComK overexpression and verified with colony PCR and Sanger sequencing (Quintara Biosciences).

### Cell harvesting and extraction

For all exponential phase steady-state measurements of *aprE* and *gfp* reporters, cells were grown from overnight starter cultures in 180 mL LB media with selective antibiotic and an inducer (20 ng/ml aTc final concentration) for at least 10 population doublings prior to reaching OD600 0.1-0.2, at which point cells were harvested by the addition of ice-cold phenol-ethanol stop solution, pelleting at 4 °C and 4000 g for 10 minutes, and storage at −80 °C until nucleic acid extraction. For steady-state MPRAs, at least 6 cell pellets were harvested per library (3 replicates for RNA and 3 replicates for gDNA; Figure S2A).

For exponential phase response time measurements of *aprE* reporters, cells were grown from overnight starter cultures in 500 ml LB medium (in 2800 mL Pyrex No. 4420 flasks) but the aTc was not added until OD 600 0.02-0.05. Cells were harvested, as described above, immediately prior to the addition of aTc (t_0_) and for the subsequent ∼60 minutes at specified time intervals.

For *ctc* reporters, single colonies of each strain were grown to OD 600 0.5 in 2 ml MOPS rich media for *B. subtilis* (Supplementary Table S4) and back diluted to OD 600 0.0002 in 15 ml media. This chemically defined rich media was used for its lack of autofluorescence, allowing for sensitive measurements of mNeonGreen level. Upon reaching an OD of 0.1-0.2 (9-10 doublings in exponential growth), 8 ml of culture harvested as described above. Spectinomycin was added to the remaining culture to a final concentration of 1 mg/mL and four technical replicates of each culture were plated in a 96-well plate for the fluorescence assay described below.

For RNA extractions, cell pellets were treated with 100 µl 10 mg/ml lysozyme in TRIS pH 8 for 5 minutes and total RNA was extracted using the RNeasy Plus Kit (Qiagen) after passing through the gDNA eliminator column. For gDNA extractions, cell pellets were handled with the gram-positive specifications of the Wizard Genomic DNA Purification Kit (Promega) and the purified gDNA was rehydrated in 100 μl at 65 °C for 1 hour. The gDNA was further purified using a 2X ratio of SPRI beads and resuspended in 100 μl water. Additional information and library-specific details are available in the extended methods sections.

### MPRA quantification

For each steady-state MPRA, six sequencing libraries were prepared: three to quantify the 8mers in the purified RNA replicates and three to quantify 8mers in the purified gDNA replicates (Figure S2A). For the cDNA libraries, 4 µg of purified RNA was reverse transcribed for 45 minutes at 50 °C in a 20 μl reaction using SuperScript III RT (Invitrogen) after incubating with 2 μl of 100 μM random hexamers (Thermo Fisher Scientific) at 65 °C for 5 minutes. RNA was hydrolyzed with 2.5 μl 1 M NaOH for 15 minutes at 95 °C and neutralized with 2.5μl HCl. The cDNA was further purified with a 0.7X ratio of SPRI beads and resuspended in 25 μl water.

A 1st PCR was performed for 2 cycles to add unique molecular identifiers (UMIs) using oligos annealing as far as possible from the random 8mer to reduce bias (Figure S3 and described in ‘MPRA optimization’ below). For each 1st PCR, approximately 1 μg (∼10 μl) of the generated cDNA or 1 μg (∼10 μl) of the purified gDNA was used in a 150 μl PCR with Phusion HF DNA Polymerase (NEB) and corresponding oligos (Supplementary Table S3). The 1st PCR amplicon was purified twice with a 1.1X ratio of SPRI beads and resuspended in 100 μl water.

Cycle-course 2nd PCR tests were performed across 8-16 cycles for a subset of samples using a subset of each 1st PCR with oligos corresponding to the next-generation sequencing platform used (Element Biosciences AVITI, Illumina NextSeq, or Singular Genomics; Supplementary Table S3) in 25 μl reactions with Phusion HF DNA Polymerase (NEB). Final library prep 2nd PCRs were performed at the identified optimal cycle number using unique indices and the remaining volume of each 1st PCRs across multiple 25 μl PCRs set up similarly to the cycle-course tests. 2nd PCR amplicons were purified with a 1X ratio of SPRI beads and resuspended in 30 μl water prior to validating on the Advanced Fragment Analyzer (AATI) and high-throughput sequencing. Additional information and library-specific details are available in the extended methods sections.Next-generation sequencing of RNA and gDNA counts (Figure S2A-B) were calculated after setting a Q30 sequencing quality score threshold for each position within the 8mer. The corresponding 15-nt UMIs allowed identification of any jackpotting during PCR amplification. Counts from gDNA replicates were used to determine the noise-threshold of at least 50 combined gDNA counts which was used to remove any lowly abundant 8mers in the library from subsequent analysis (Figure S2C). PyDESeq2 (Muzellec et al., 2023) with dispersion and shrinkage was used on 3 replicate RNA counts and 3 replicate gDNA counts to compute the fold-change for each 8mer (reported as its frequency_RNA_ / frequency_DNA_) and their corresponding adjusted *p-*value was computed using the Benjamini and Hochberg method (displayed only for 8mers above the noise threshold). Corresponding sequencing files and code will be publicly available. Additional information and library-specific details are available in the extended methods sections.

### MPRA optimization

The library preparation protocol (described in ‘MPRA quantification’ above) was converged upon by incrementally optimizing the reverse transcription and 1st PCR set-up to minimize measurement biases generated at those steps, while still fitting the length confines of a 75-bp paired-end short-read sequencing run. The *aprE* +997-nt site library was used for these optimizations. Use of a gene-specific RT primer that binds immediately downstream of the random 8mer site revealed 959 8mers with a log2 freq_RNA_/freq_DNA_ < −0.5 and an adjusted *p* < 0.05 (Figure S3A). The corresponding gDNA libraries were prepared by using the gene-specific RT primer as the reverse primer for the 1st PCR and purified double-stranded genomic DNA as the template. Levenshtein edit-distance based clustering of these statistically significant repressive variants revealed several clusters (Figure S3B). The largest of these clusters contained motifs with sequence complementary to the gene-specific RT primer’s binding site, suggesting that 8mers that occluded RT primer binding lowered the measured freq_RNA_/freq_DNA_ by inhibiting cDNA synthesis (Figure S3B, inset).

Use of proximal PCR primers (with a forward primer binding sites 5 nucleotides upstream and reverse primer binding site immediately downstream of the 8mer site) after reverse transcribing with random hexameric RT primers revealed 311 8mers with a log2 freq_RNA_/freq_DNA_ < −0.5 and an adjusted *p* < 0.05 (Figure S3C). Both the cDNA and gDNA libraries were prepared using the same set of proximal PCR primers for the 1st PCR. Levenshtein edit-distance based clustering of these 311 statistically significant repressive variants revealed distinct clusters (Figure S3D), the largest of which contained motifs with sequence complementary to the proximal PCR primers’ binding sites, suggesting that they were experimental artifacts. These variants with low log2 freq_RNA_/freq_DNA_ were further corroborated as false-positives using high-throughput half-life measurements (see ‘Cell harvesting and extraction’) of the +997-nt site library. Sequencing reads for the RNA level of each 8mer at each time-point were normalized to the level of the *sigY* transcript, a reference gene whose expression does not change during the aTc induction time-course. Only 8mers measured above the noise-threshold (more than 200 sequencing read counts at the 60 min time-point) were used to fit curves (see ‘mRNA half-life measurement’). The cumulative probability distribution of the half-lives of these low log2 freq_RNA_/freq_DNA_ variants was not shifted to the left relative to the distribution of all 8mers (Figure S3E), indicating that these variants did not lower molecular half-lives but lowered the freq_RNA_/freq_DNA_ due to the bias in experimental measurement.

The bias was partially corrected by using a distal downstream PCR primer (with a binding site 76 nt downstream of the 8mer site) after reverse transcription with a random hexameric RT primer. Both the cDNA and gDNA libraries were prepared using a proximal forward primer and a distal downstream primer for the 1st PCR. This revealed 147 8mers with a log2 freq_RNA_/freq_DNA_ < −0.5 and an adjusted *p* < 0.05 (Figure S3F). Levenshtein edit-distance based clustering of the 147 statistically significant repressive variants revealed no 8mers complementary to the distal downstream PCR primer’s binding site. However, the largest cluster contained a motif with sequence complementary to the upstream forward primer’s binding site (Figure S3G and inset).

The bias was further corrected by using a distal upstream PCR primer (with a binding site 33 nt upstream of the 8mer site) and a distal downstream PCR primer (with a binding site 76 nt downstream of the 8mer site) after reverse transcription with a random hexameric RT primer. Both the cDNA and gDNA libraries were prepared using a distal forward primer and a distal downstream primer for the 1st PCR. This revealed only 39 8mers with a log2 freq_RNA_/freq_DNA_ < −0.5 and an adjusted *p* < 0.05 (Figure 3SH), notably fewer than those previously observed at this +997-nt site and without the previously observed false-positives. The largest cluster observed, from Levenshtein edit-distance based clustering of the 39 statistically significant repressive variants (Figure S3I), contained a motif with sequence complementary to a loop at the base of the 3′ end stabilizing hairpin. This small degradation effect was confirmed by the high-throughput half-life measurements which demonstrated a slight decrease in the half-life of cluster D sequences (Figure S3J). Together, these iterative optimizations revealed that amplifying distally after random hexameric RT priming minimized RT and PCR biases in MPRA barcode quantification and all MPRAs reported in this study (for *aprE* and *gfp*) were carried out with this scheme of distal amplification. Additional information and library-specific details are available in the extended methods sections.

### Anti-RBS and *k*-mer analysis

Anti-RBS 8mers were defined as those that have a free energy of binding to the RBS less than −8 kcal/mol. The minimum free energy was calculated by ViennaRNA’s RNAcofold using the 40-nt RBS (5′-AAAGGAGAGGGUAAAGA<u>GUG</u>AGAAGCAAAAAAUUGUGGAU-3′, spanning from −17 to +23 relative to the underlined GUG start codon of the reporter). Distributions of freq_RNA_/freq_DNA_ are only shown for anti-RBS 8mers above noise-threshold in the corresponding library.

For each of *k* = 4, 5, 6, 7, and 8, all possible *k*-mers were generated. Then, for each *k*-mer, its instances in the +467-nt random 8mer library were identified after accounting for *k*-1 nucleotides of constant sequence on either side. The freq_RNA_/freq_DNA_ was only tabulated for instances where the variant in the +467-nt random 8mer library was measured above the noise-threshold. Then, the mean freq_RNA_/freq_DNA_ for each *k*-mer was computed as the average of its tabulated freq_RNA_/freq_DNA_ ratios.

Levenshtein edit-distances were calculated (using the distance.levenshtein method of the Distance package for Python) amongst the 500 lowest ranked *k*-mers of length 8 from the *k-*mer analysis of the +467-nt library described above. Hierarchical clustering of the Levenshtein edit-distance matrix was performed using the seaborn.clustermap method of the Seaborn package for Python, and clusters were extracted from the linkage map using the hierarchy.fcluster method of the SciPy package for Python. Sequence motifs after edit-distance clustering were identified from each cluster using Multiple Em for Motif Elicitation (MEME) in the classic mode using a zero or <u>o</u>ne <u>o</u>ccurrence <u>p</u>er <u>s</u>equence (zoops) site distribution, a uniform background set, and searching for motifs between 2 and 8 wide (inclusive). Statistically significant motifs with the lowest E-values from MEME outputs were plotted in Python using Logomaker (Tareen and Kinney, 2020).

Reading frame analysis for each of the lowest ranked *k*-mers of length 4, 5, and 6 was performed by identifying instances in the +467-nt random 8mer library of that *k*-mer at specific positions and tabulating the freq_RNA_/freq_DNA_ ratios of those instances. The distributions of their tabulated freq_RNA_/freq_DNA_ ratios were plotted using the pyplot.boxplot method of the Matplotlib package in Python. Distributions of in-frame and out-of-frame instances of proline codons, pairs of proline codons, and stop codons were similarly obtained by tabulating the freq_RNA_/freq_DNA_ ratios of their instances in the +467-nt random 8mer and plotted using the using the kernel density estimate (KDE) method of the Seaborn package in Python. Details regarding the quantification and statistical analysis of the data can be found in the relevant parts of the results sections and legends. Corresponding code will be publicly available.

### RT-qPCR quantification

For qPCR measurements across all reporters, 1 µg of purified RNA was reverse transcribed in a 10 μl reaction using MuLV RT (NEB) after incubating with 1 μl of 100 μM random hexamers (Thermo Fisher Scientific) at 65 °C for 5 minutes. After reverse transcription was complete, RNA was hydrolyzed with 2 μl 1 M NaOH for 5 minutes at 95 °C and neutralized with 2 μl HCl prior to diluting in 86 μl 10 mM Tris pH 8. 2 μl of this 1X (or a further 1:10X diluted) cDNA used as template in a 384-well qPCR with reporter-specific oligos (Supplementary Table S3) and 2X KAPA SYBR Green Master Mix (Sigma-Alrich) on a Roche LightCycler 480 Real-Time PCR machine. The *B. subtilis* gene *gyrA* was used as a reference gene for computing mRNA levels. C_t_ values for each locus were calculated by averaging the C_t_ values of technical replicates and corrected for primer efficiency. The relative mRNA values for each reporter construct were obtained by raising 2 to the power of C_t_reference_-C_t_reporter_ and further normalized as specified in the corresponding results.

### mRNA half-life measurement

Cells from exponential-phase time-courses were harvested and extracted for RT-qPCR or MPRA quantification as described elsewhere and data were fit using the following formulism (Alon, 2007). A mathematical model of the rates underlying the dynamics of mRNA molecules whereby the mRNA is transcribed from DNA by a production rate (beta, β) and the mRNA decays with a decay rate (alpha, α) can be given written as:

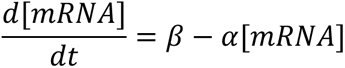

Here, the decay rate is a combination of the molecular degradation rate and the rate of molecular dilution during cell division. The steady-state level is dependent upon both β and α as follows:

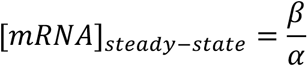

Upon induction, the dynamic equation results in an approach to steady-state described by:

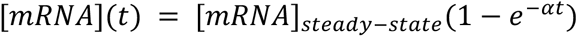

The above is used to fit the induction time-course data to obtain a value for α. Further, solving for the time to reach half of the steady-state level, we find that the response time, t_1/2_, is given by:

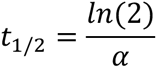

This response time upon inducing production is the same as the “half-life” obtained from stopping production and observing exponential decay of the concentration of mRNA. Notably, the response time for both increase and decrease in mRNA levels is not dependent on β and is only dependent on α. Specifically, the dependence on α is inverse, i.e., the larger the decay rate, the shorter the response time.

### DMS-MaPseq and mRNA secondary structure analysis

DMS treatment protocol was adapted from previous publications (McCormick et al., 2021; Zubradt et al., 2017; Burkhardt et al., 2017). Briefly, overnight cultures were diluted 10,000-fold into 50 ml of LB media with the selected antibiotic and inducer as described above. At OD 600 0.2, the culture was split into two 15 ml aliquots. For each strain, one 15 ml culture was incubated with 750 µl of dimethyl sulfate (DMS, ∼5% final concentration) at 37 °C in a shaking water bath for 2 minutes. The reaction was stopped by adding 30 ml of ice cold 30% β-mercaptoethanol in 1X PBS pH 7.4 and immediately pelleted by spinning at 4000 g for 8 minutes at 4 °C. Pellets were then washed with 8 ml of ice cold 30% β-mercaptoethanol and pelleted once more before being frozen at −80 °C.

Cell pellets were centrifuged for 3 minutes at 8000 g to thaw and remaining supernatant was removed. Pellets were then treated with 100 µl 10 mg/ml lysozyme in TRIS pH 8 for 5 minutes in 400 µl RNA lysis buffer (10 mM EDTA, 50 mM sodium acetate). Total RNA was extracted using hot acid-phenol:chloroform followed by isopropanol precipitation.

For sequencing amplicon preparation of the *aprE* reporter, samples were treated with TURBO DNase followed by RNA Clean & Concentrate −5 (Invitrogen, Zymo). Reverse transcription was performed at 55 °C for 10 minutes using Induro reverse transcriptase (NEB) and a gene-specific RT primer. RNA was hydrolyzed with 4 μl 1 M NaOH for 5 minutes at 95 °C and neutralized with 4 μl HCl. PCR amplification of the RT product was performed using Phusion High-Fidelity DNA polymerase, using the RT reaction as the template and the RT primer as the reverse primer for 11-13 cycles. The resulting 163 nt product was purified using 1.8X SPRI bead selection and resuspended in a final 100 µl of water. Adapter sequences were added using NEBNext Ultra™ II DNA PCR-free Library Prep Kit for Illumina as per manufacturer’s recommendation. A second PCR was performed using Phusion High-Fidelity DNA polymerase and indexing oligos corresponding to Illumina NextSeq (Supplementary Table S3) for 4-7 cycles and purified using 1.8X SPRI beads. The libraries were sequenced on an Illumina MiSeq NANO (2×150 nt reads).

For sequencing amplicon preparation of the *ctc* reporter, RNA samples were similarly treated with TURBO DNase and RNA Clean & Concentrate −5 (Invitrogen, Zymo). Reverse transcription was performed at 60 °C for 30 minutes using Induro reverse transcriptase (NEB) and a gene-specific RT primer. RNA was hydrolyzed with 4 μl 1 M NaOH for 5 minutes at 95 °C and neutralized with 4 μl HCl. PCR amplification of the RT product was performed using Q5 DNA polymerase using the RT reaction as the template and the RT primer as the reverse primer for 15-20 cycles, until detectable on a TBE polyacrylamide gel. The resulting 199 nt amplicon was purified using 1.8X SPRI beads and adapter sequences were added using NEBNext Ultra™ II DNA PCR-free Library Prep Kit for Illumina as per manufacturer’s recommendation. A second PCR was performed using Q5 DNA polymerase and indexing oligos corresponding to Illumina NextSeq (Supplementary Table S3) for 6-9 cycles and purified twice, with SPRI beads, first at a 1X then a 1.8X concentration. The libraries were sequenced on an Illumina MiSeq i100 instrument (2×150 nt reads) with a 50% PhiX spike in to increase sample complexity.

To determine DMS reactivity, FASTQ files were analyzed using the SEISMIC pipeline (Allan et al., 2024) with default settings. Primer annealing sites were masked out using -P. For each sample, signal was normalized using seismic graph profile --use-ratio --quantile 0.95. Computationally generated MFE structure, constrained with DMS reactivity data for the regions for which it was measured, was generated for the full transcript and visualized using RNAfold 2.6.3 (Gruber et al., 2008).

mRNA secondary structure analysis for stabilizing 8mers was performed using ViennaRNA’s RNAfold by extracting the total base-pairing probability of every nucleotide in the full-length transcript. The ratio of base-pairing probability of each nucleotide in the SD was computed by dividing the median of 100 randomly sampled stabilizing 8mers (freq_RNA_/freq_DNA_ > 1.75) from the −25-nt 5′ UTR MPRA to the median of 100 randomly sampled invariant 8mers (0.75 < freq_RNA_/freq_DNA_ < 0) introduced at that site.

### Fluorescence quantification

To quantify mNeonGreen protein levels, measurements of absorbance (600 nm) and fluorescence intensity (EX 485/20 nm, EM 510/20 nm) were made in a BioTek Synergy H1 microplate reader. Upon reaching an OD of 0.1-0.2, cultures were treated with spectinomycin (final concentration of 1 mg/mL) and four technical replicates of each culture were plated in a 96-well (see Cell harvesting and extraction). Fluorescence values and OD values were corrected by subtracting the background signal from wells containing media only, and relative fluorescence of each strain was obtained by dividing the corrected fluorescence value by the corrected OD value. Cellular autofluorescence was calculated by performing these measurements on wildtype *B. subtilis* cells not expressing any fluorescent protein, and this value was subtracted from all fluorescent strains.

### *In vitro* translation and Western blot

Reporters were cloned into a backbone containing a T7 RNA polymerase promoter and terminator. RNA was then *in vitro* transcribed from 1 µg plasmid for 90 minutes at 37 °C using T7 RNA polymerase (NEB). Remaining DNA was removed with 1 µl of TURBO DNase (Invitrogen) and RNA was purified using Zymo RNA Clean & Concentrate −25. RNA was then quantified before being melted and refolded either with or without antisense DNA oligos (IDT). The reaction was suspended in refolding buffer (2 mM MgCl_2_, 100 mM NaCl, 10 mM TRIS pH 8), heated to 94 °C and slow cooled to room temperature. 600 ng of RNA ± ASO was loaded into a 10 µl PURExpress *in vitro* translation reaction and incubated at 37 °C for 3 hours. RNA concentration was also verified on an Advanced Fragment Analyzer (AATI) to confirm that minimal degradation occurred during melting and refolding. Reactions were stored at −20 °C after completion.

To perform the Western blot, 4 µl of the PURExpress reaction was mixed with 4 µl of 4X Fluorescent Compatible Sample Buffer with 5% β-mercaptoethanol and denatured for 3 minutes at 70 °C (Thermo Fisher). Protein was then run on a NuPage 4-12% Bis-Tris gel for 40 minutes and then dry transferred with an iBlot for 5 minutes (Invitrogen). Membrane was then placed in 1X Blocker™ FL Fluorescent Blocking Buffer for 45 minutes before incubating with monoclonal C163 α-GFP primary antibody overnight at 4 °C (Thermo Fisher). The membrane was then washed 3 times in TBST (TRIS pH 7.6, 0.01% Triton X-100) and incubated in blocking buffer with α-Mouse secondary antibody conjugated to Alexa Fluor™ Plus 647 fluorophore for 30 minutes at room temperature. After washing 3 more times in TBST, the membrane fluorescence was imaged on a Typhoon FLA 9500 imager.

### aSD-like depletion analysis

aSD-like 8mers were defined as those that have a free energy of binding to the consensus SD less than −8 kcal/mol. The minimum free energy was determined by ViennaRNA’s RNAcofold using the consensus 9-nt bacterial SD (5′-AGGAGGUGA-3’). This procedure yields a set of 882 aSD-like 8mers. Equivalently sized set of 882 control 8mers were obtained by randomly sampling from all possible 8mers. Bacterial coding sequences were obtained from annotations of their reference genomes (NC_000964.3, ASM584v2, and CP001340.1 for *B. subtilis* 168, *E. coli* K-12 MG1655, and *C. crescentus* NA1000, respectively.) Annotated CDSs of lengths non-divisible by 3 were excluded, yielding sets of 4,218 *B. subtilis* CDSs, 4,146 *E. coli* CDSs, 374 *C. crescentus* leaderless CDSs (leaderless identified from Dataset S1 of Schrader et al., 2014), and 418 *B. subtilis* simple TU CDSs (simple TUs identified from Table S5 of Herzel et al., 2022).

The total number of occurrences of each aSD-like 8mer in a set of CDSs was determined by finding exact sequence matches and tabulating the number of its occurrences and their observed positions among all potential CDSs. The frequency of an aSD-like 8mer was calculated by dividing its total number of occurrences by the total number of CDSs in that set. The same was repeated for each 8mer from the control set and their respective distributions were plotted using the KDE method of the Seaborn package in Python. Statistical tests were performed with SciPy package in Python.

For specific bin sizes, positional frequencies in a bin were obtained by dividing the total number of 8mers observed in that bin by the number of CDSs whose length is longer than that bin.

Shuffling of CDSs was performed by keeping the start and stop codons intact. For nucleotide shuffling, the random.shuffle method of the NumPy package in Python was used to scramble the positions of nucleotides in each CDS while maintaining its nucleotide frequency. For triplet shuffling, the CDS was split into chunks of 3 nt before similarly scrambling the positions of triplets while maintaining the triplet frequency of each CDS. Codon shuffling was performed per the CodonShuffle method (Katz and Burge, 2003) which preserves the encoded protein sequence and the same codon usage of each native CDS. This is done by randomly permuting the set of codons used within each CDS while maintaining the order of the encoded amino acids observed in that CDS. The shuffling was repeated 20 times independently for each CDS in each set of CDSs. The frequencies of the 882 aSD-like 8mers was computed in each of the 20 shuffled sets and the mean frequency of each aSD-like 8mer was computed. The distribution of aSD-like frequencies was plotted using the KDE method of the Seaborn package in Python. Statistical tests were performed with SciPy package in Python. Details regarding the quantification and statistical analysis of the data can be found in the relevant parts of the results sections and legends. Corresponding code will be publicly available.

## SUPPLEMENTAL FIGURES

**Figure S1.**
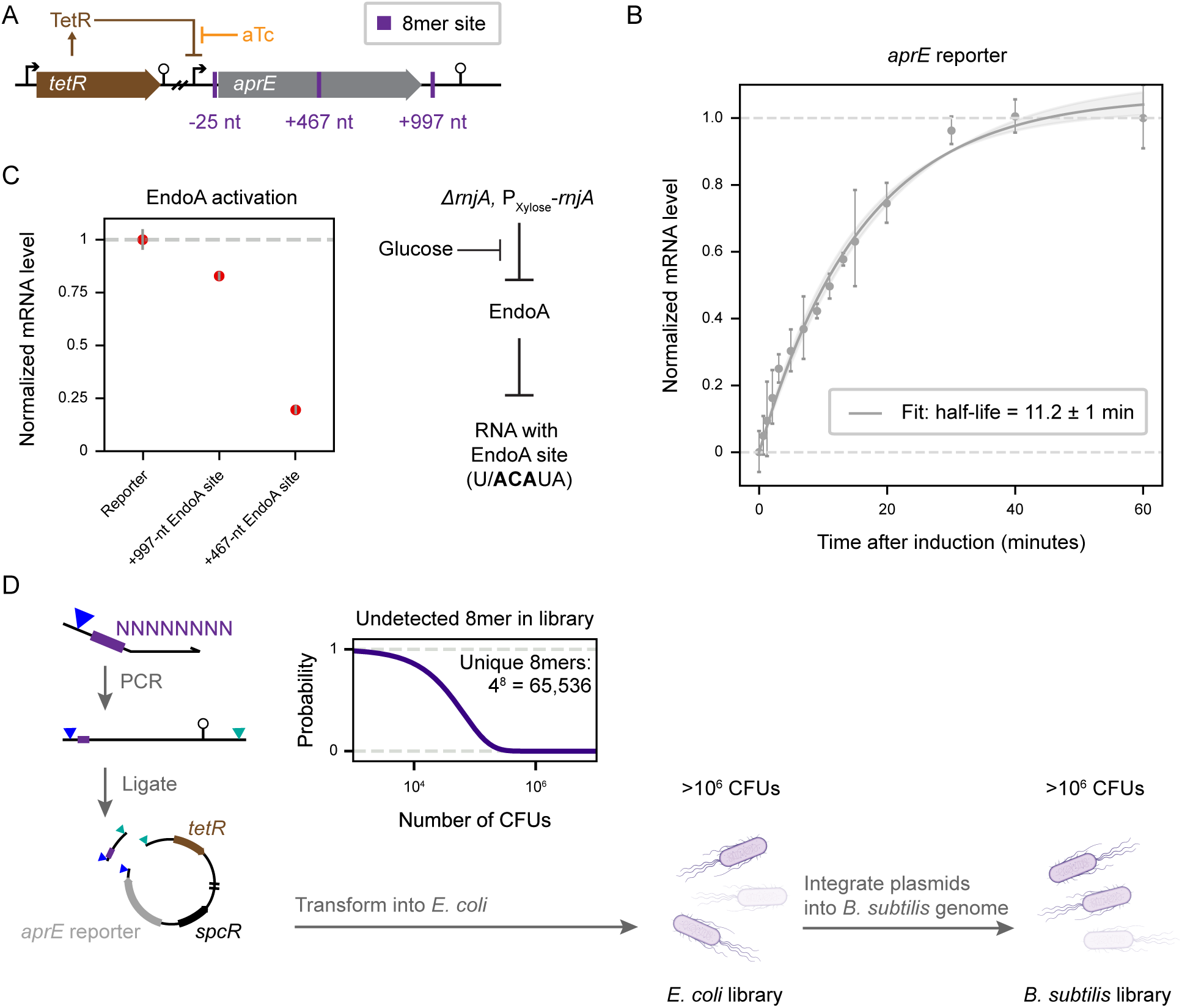
The inducible *aprE* reporter produces a stable mRNA transcript and can report on mRNA degradation. (**A**) Schematic of the aTc inducible *aprE*-derived reporter (modified from the native gene as described in Methods) with the −25-nt, +467-nt, and +997-nt sites highlighted. (**B**) Half-life measurement from an induction curve of the reporter measured by RT-qPCR. Error bars represent propagated standard error from 3 technical replicates and shaded region represents the 95% confidence interval of the fit. (**C**) Schematic for ectopic activation of the EndoA nuclease in this strain of *B. subtilis* by depletion of RNaseJ1. Normalized mRNA levels of reporters with EndoA cleavage site in the +997-nt site or at the +467-nt site show destabilizing effect of EndoA activity but is more pronounced in the CDS, likely due to the high prevalence of secondary structure around the +997-nt site. (**D**) Schematic of library construction in the reporter to obtain all possible 8mer variants at a specific site. Restriction sites marked as triangles. Inset: Simulation to predict number of transformants needed at every step of the protocol to maintain library diversity.

**Figure S2.**
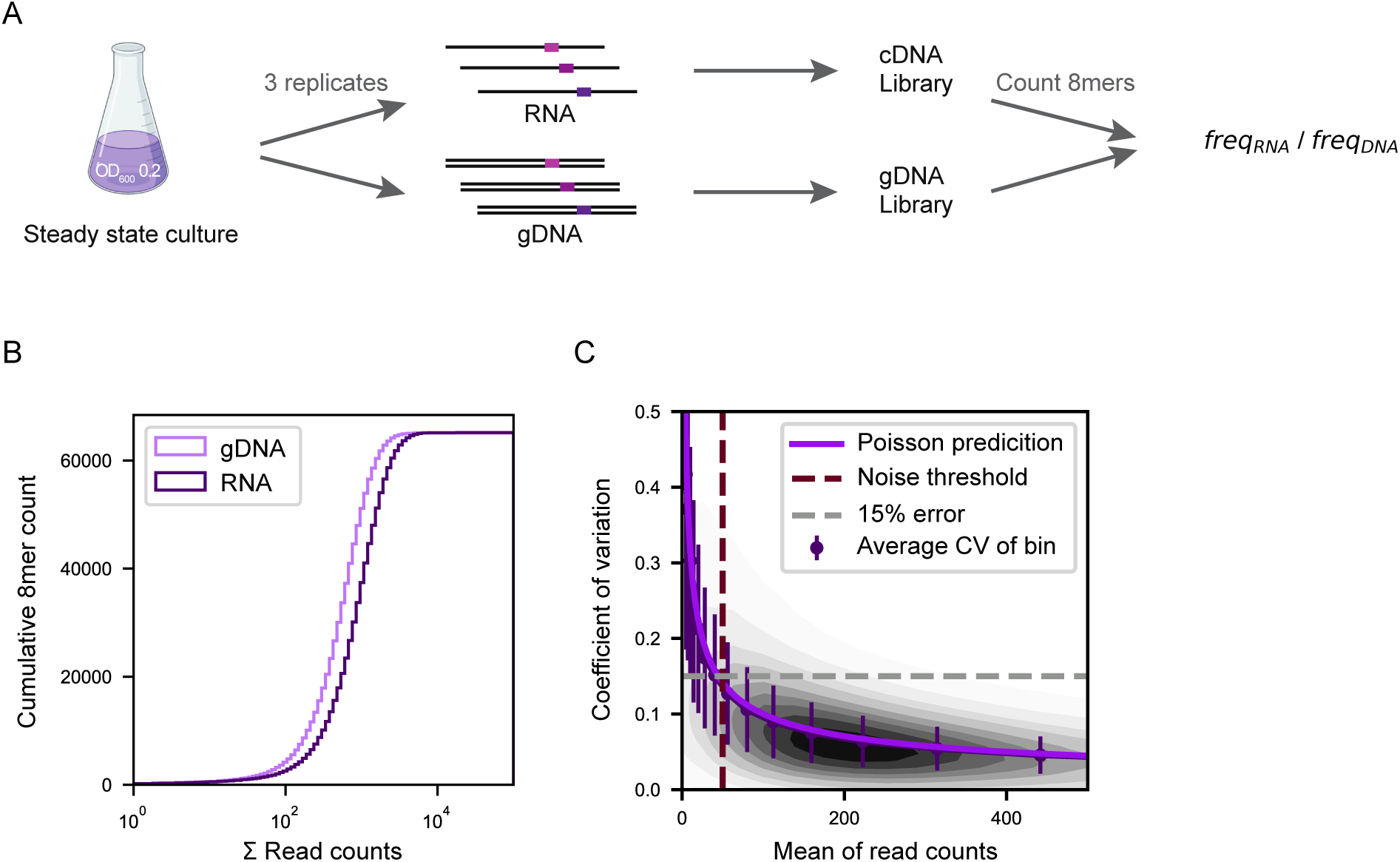
Overview of RNA-seq based MPRA measurement. (**A**) RNA-seq based workflow for obtaining steady-state mRNA levels of the 8mer variants. (**B**) Plot showing the typical distribution of total RNA and gDNA sequencing read counts obtained from the above workflow. Total gDNA counts of 67M and total RNA counts of 117M were obtained for the library shown. (**C**) Assessment of noise in read counts performed by leveraging technical replicates (n = 3) and counting UMIs of each 8mer variant, compared to Poisson predictions. 50 sequencing read counts at the gDNA level was set as the threshold for all subsequent analysis (Methods).

**Figure S3.**
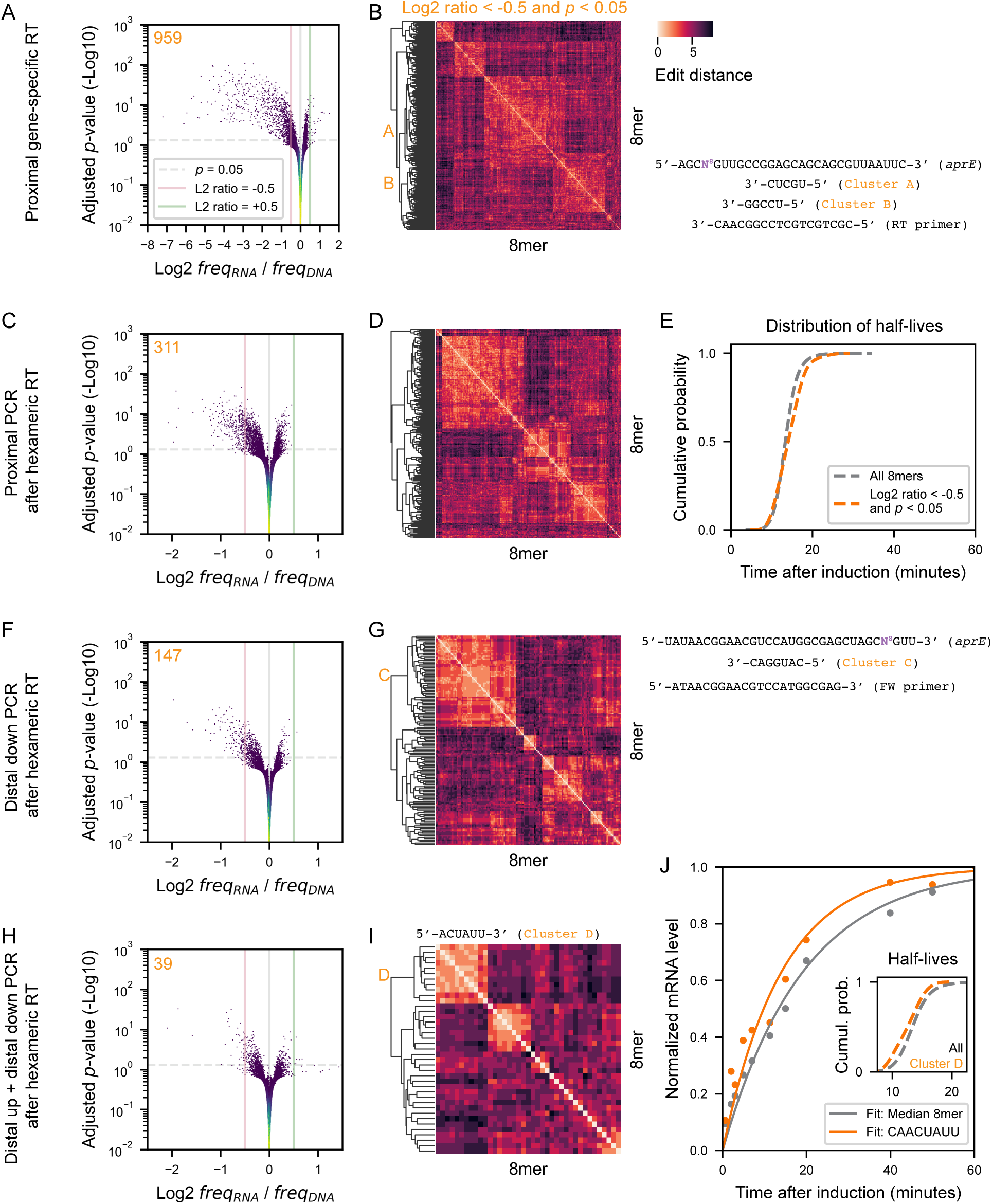
Amplifying distally after random hexameric RT primer minimizes biases in MPRA barcode quantification. (**A**) Using a proximal gene-specific RT primer. Plot of the steady-state level of mRNA variants with 8mers in the +997-nt site of the *aprE* reporter shown as the log2 freq_RNA_/freq_DNA_ (L2 ratio) normalized around the median in the library and their corresponding Benjamini-Hochberg adjusted *p*-value. Colored according to kernel density estimation of number of variants per pixel. 959 8mers with a log2 freq_RNA_/freq_DNA_ < −0.5 and an adjusted *p* < 0.05. (**B**) Levenshtein edit-distance based clustering of the 959 statistically significant repressive variants from **A**. Inset: Schematic of the +997-nt random 8mer site (N^8^) of the *aprE* reporter showing the binding sites of the RT primer and the statistically significant motifs identified from their corresponding clusters using motif enrichment analysis. 416 8mers in cluster A and 208 in cluster B. (**C-D**) Same as **A-B** from using proximal PCR primers after random hexameric RT primer. 311 8mers with a log2 freq_RNA_/freq_DNA_ < −0.5 and an adjusted *p* < 0.05. (**E**) Cumulative probability distribution of the half-lives of 8mers from **C** with log2 freq_RNA_/freq_DNA_ < −0.5 and an adjusted *p* < 0.05 compared to distribution of half-lives of all 8mers in the +997-nt library. (**F-G**) Same as **A-B** from using a distal downstream PCR primer after random hexameric RT primer. 147 8mers with a log2 freq_RNA_/freq_DNA_ < −0.5 and an adjusted *p* < 0.05. Inset: Schematic of the +997-nt random 8mer site (N^8^) of the *aprE* reporter showing the binding sites of the proximal forward (FW) 1st PCR primer and the statistically significant motif identified from the largest cluster using motif enrichment analysis. (**H-I**) Same as **A-B** from using distal downstream and distal upstream PCR primers after random hexameric RT primer. 39 8mers with a log2 freq_RNA_/freq_DNA_ < −0.5 and an adjusted *p* < 0.05. 11 8mers in cluster D. (**J**) Half-life measurements from induction curves of reporters with the median 8mer variant and an 8mer variant from cluster D. Inset: Cumulative probability distribution of the half-lives of 8mers from cluster D compared to distribution of half-lives of all 8mers in the +997-nt library.

**Figure S4.**
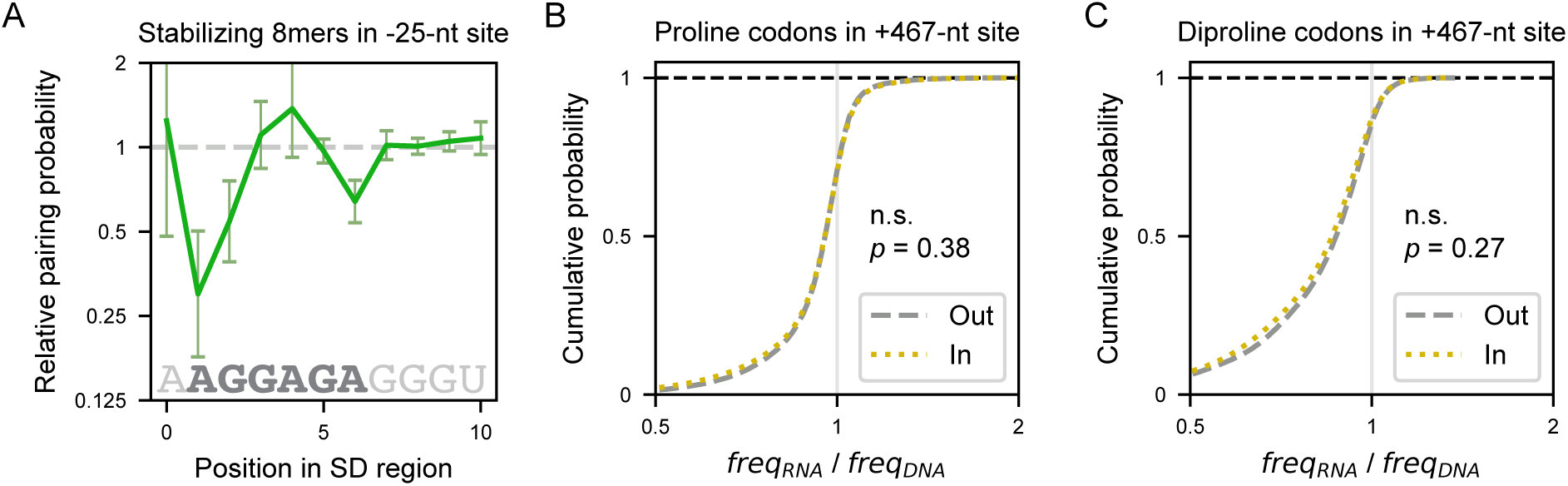
Stabilizing variants from −25-nt library increase RBS accessibility, and pairs of proline codons in the +467-nt library exhibit no reading frame dependence. (**A**) Ratio of base pairing probability of each nucleotide in the SD when comparing the median of 100 randomly sampled stabilizing 8mers from the 5′ UTR MPRA to 100 randomly sampled invariant 8mers introduced at the −25-nt site. Error bars represented the propagated standard error of the ratio. Base-pairing probabilities were obtained from *in silico* prediction of folding the full *aprE* reporter transcript using ViennaRNA’s RNAfold. Lower pairing probability represents increased accessibility of the SD to the ribosome. (**B-C**) The in-frame and out-frame effects of instances of proline codons (**B**) and pairs of proline codons (**C**) on mRNA level in the +467-nt library. *p*-values were computed from a two-sided Wilcoxon rank-sum test.

**Figure S5.**
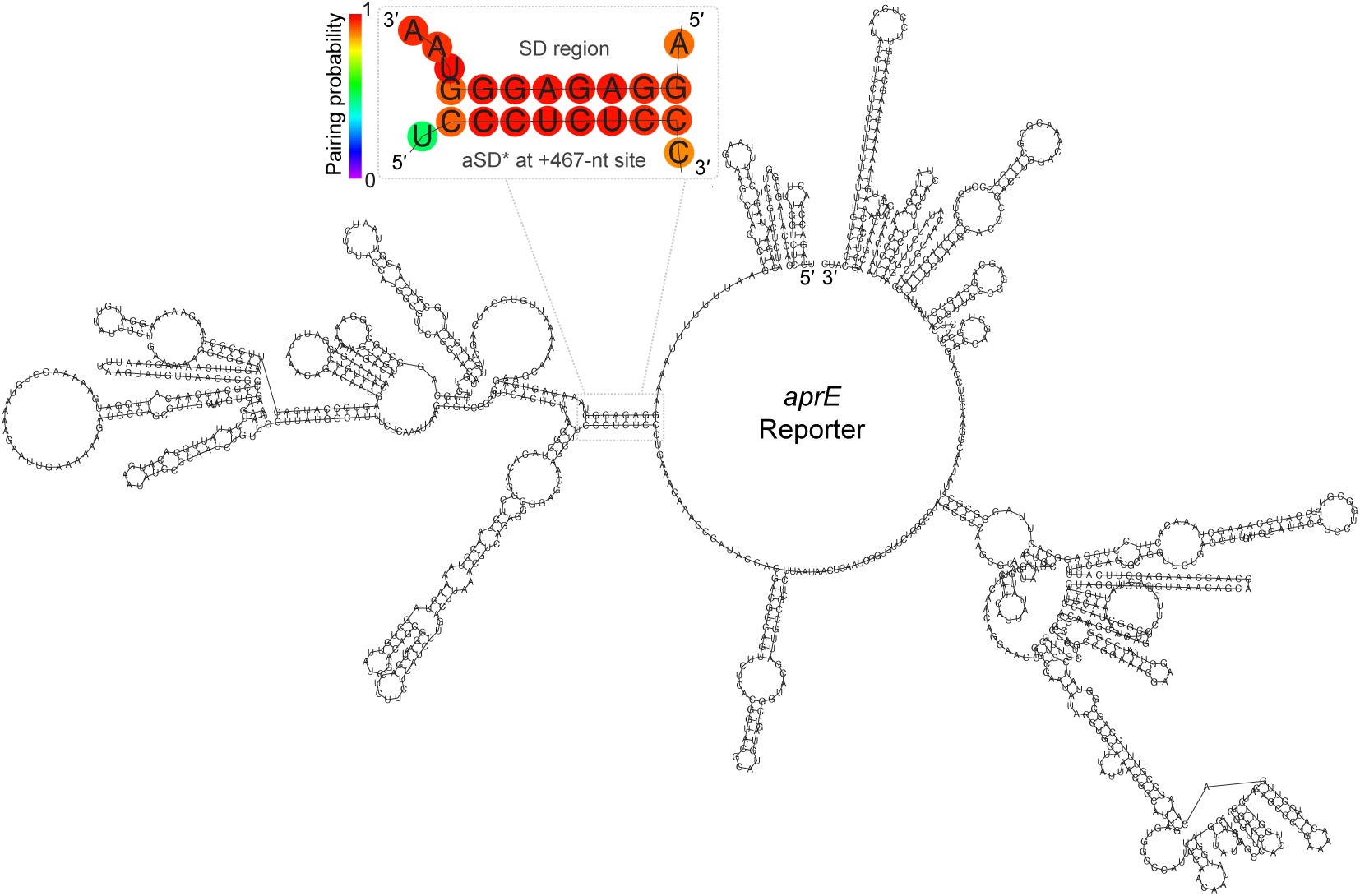
DMS reactivity constrained secondary structure of the *aprE* transcript with the aSD* 8mer at the +467-nt site shows long-range pairing with the SD region. Computationally generated MFE structure, constrained with DMS reactivity data from a 157-nt window surrounding the SD, shows that the SD region base pairs with aSD* sequence at the +467-nt site. The magnified area depicts the computationally derived base pairing probabilities. For unpaired regions, the color denotes the probability of being unpaired.

**Figure S6.**
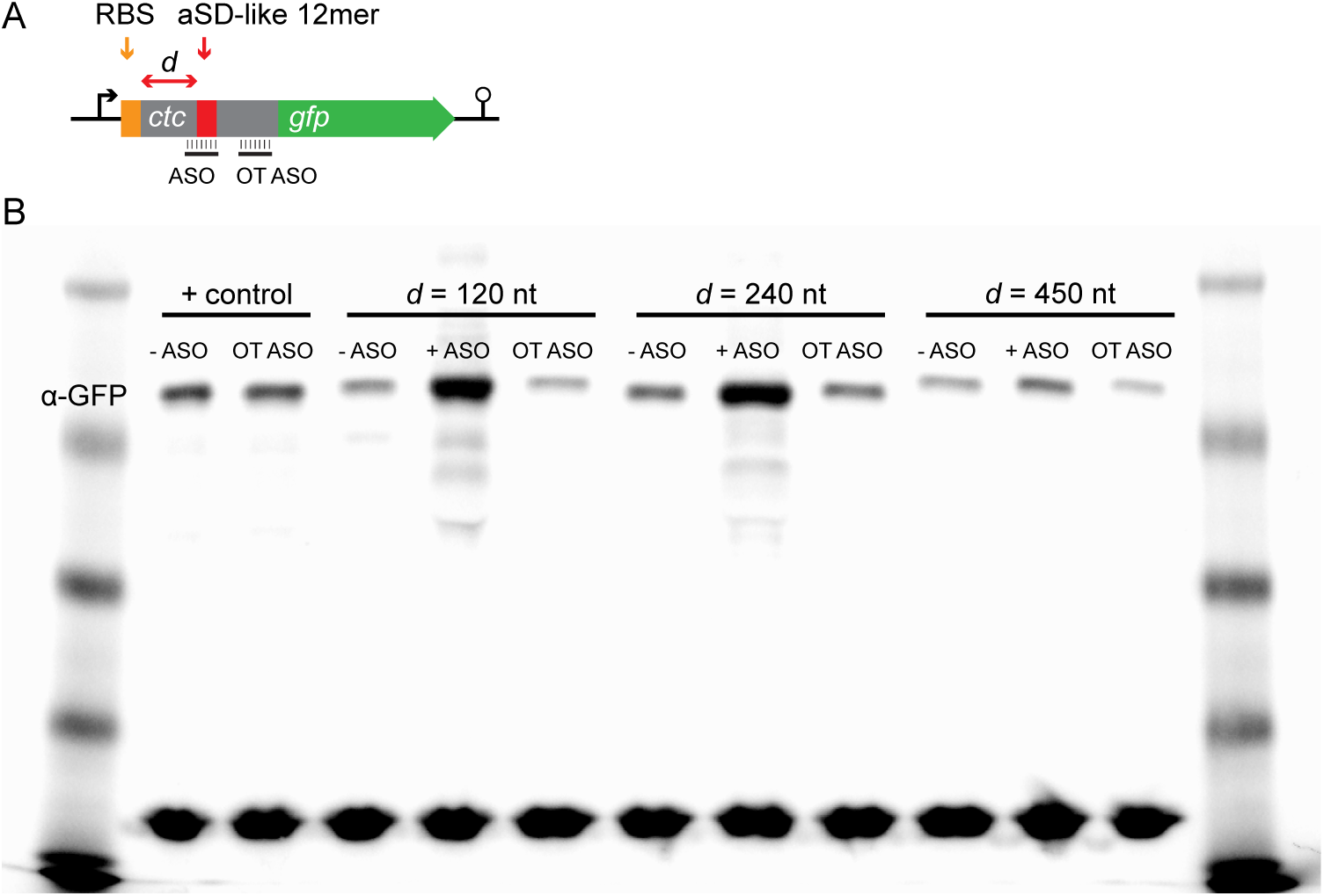
Raw anti-GFP Western blot of *in vitro* translated Ctc-GFP reporters. (**A**) Schematic of *in vitro* translated reporter (not to scale). The native *ctc* 5′ UTR and CDS are fused to *gfp* with a linker. aSD-like 12mers are substituted in at a variable distance *d* from the *ctc* start codon and antisense oligonucleotides (ASOs) bind either over the aSD-like 12mer or at an off-target site on *ctc.* ASOs ranged from 18-26 nt to achieve a T_m_ of 72-73 °C in the reaction conditions. (**B**) Anti-GFP Western blot of *in vitro* translated Ctc-GFP without an aSD-like 12mer (+ control) and with an aSD-like 12mer positioned 120, 240, or 450 nt from the *ctc* start codon ± ASO. OT ASO binds an off-target site on *ctc*.

**Figure S7.**
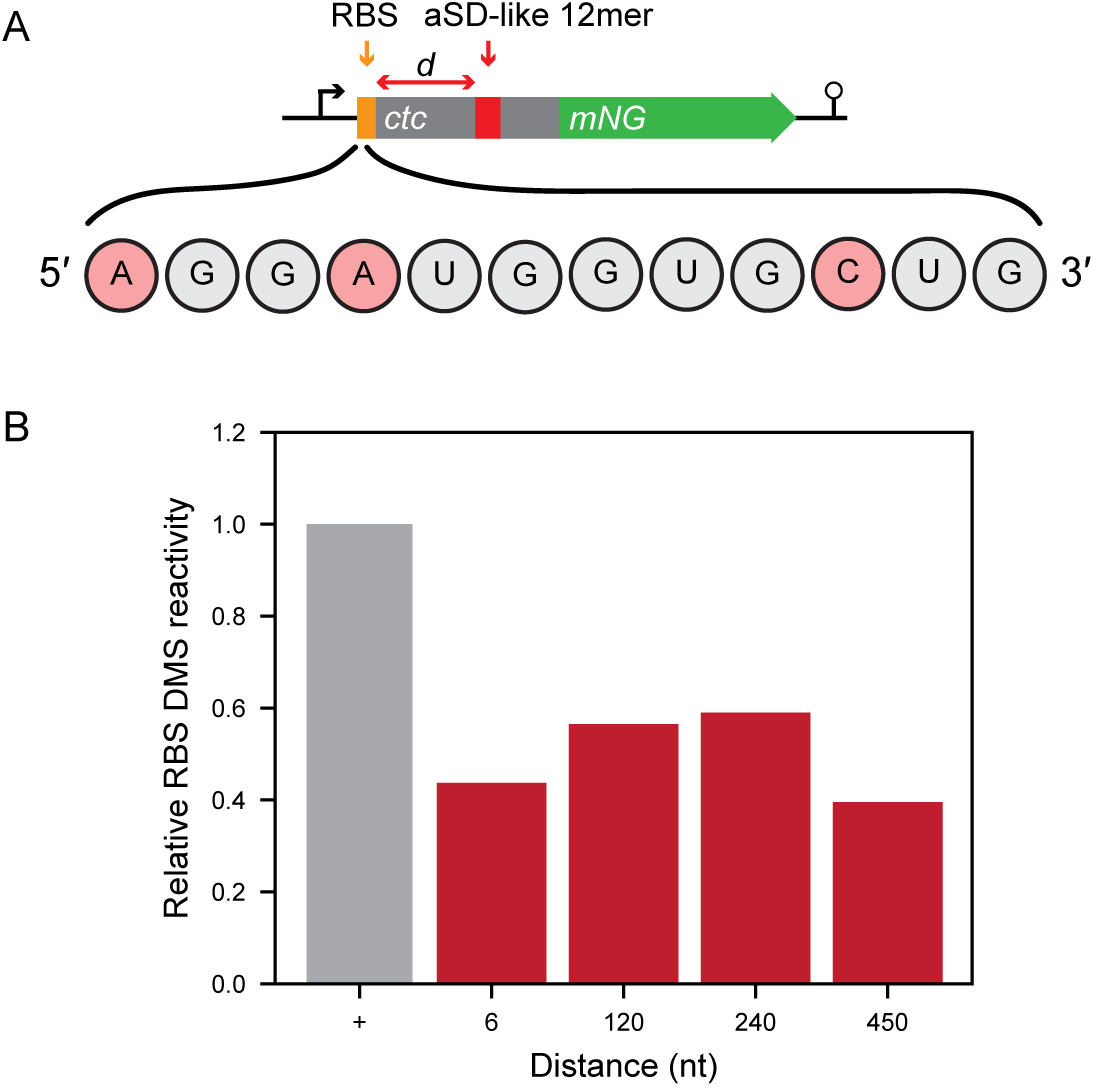
Accessibility of the *ctc* RBS to modification by DMS. (**A**) Schematic of *ctc-mNG* reporter and region nucleotides probed for DMS reactivity (not to scale). Highlighted regions include a constant RBS in the 5′ UTR and an aSD-like 12mer, complementary to the SD sequence and following 6 downstream nucleotides, that is placed in various locations along the CDS for different variants. The positive control is the native *ctc* sequence and does not contain an aSD-like mutation. Pink bases (A and C) are susceptible to modification by DMS. The 12 highlighted nucleotides comprise the section of the RBS that is targeted by the aSD-like 12mer in the different TE reporter variants. (**B**) Average DMS mutationn rate of the three RBS bases susceptible to DMS modification in *ctc* reporters with the aSD-like 12mer located 6, 120, 240, and 450 nt away from the start codon relative to the positive control (no aSD-like 12mer).

**Figure S8.**
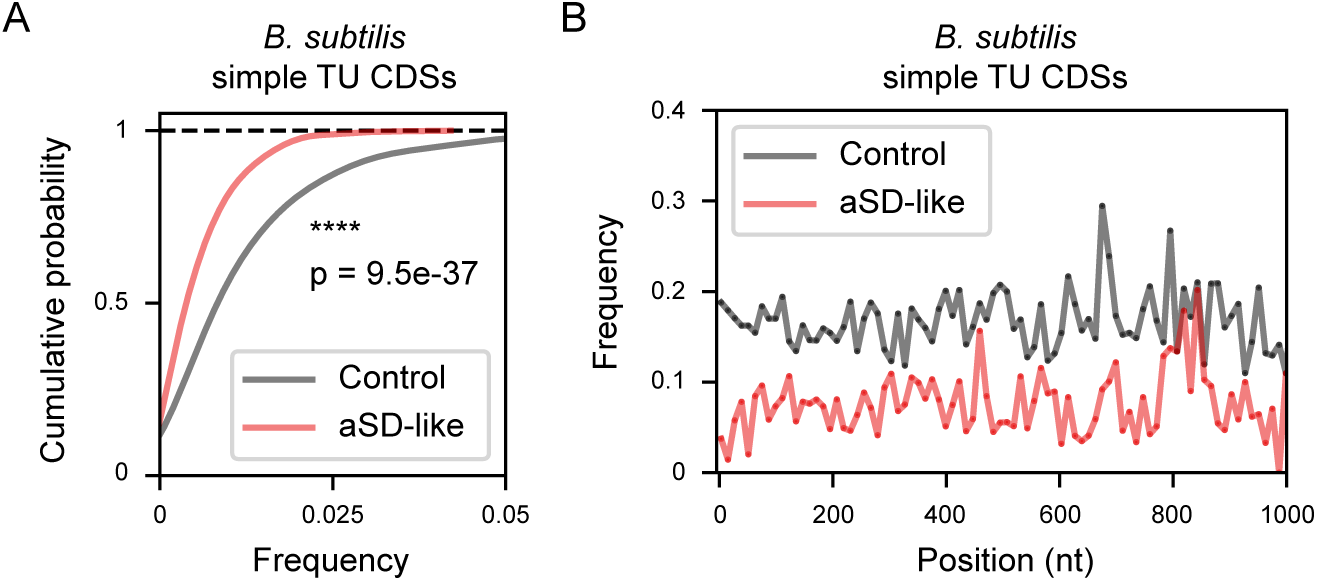
Depletion of aSD-like 8mers throughout *B. subtilis* CDSs in simple transcription units. (**A**) Frequency of aSD-like 8mers (set size = 882; free energy of binding to the consensus SD less than −8 kcal/mol) in *B. subtilis* CDSs of simple monocistronic transcription units (TUs, n = 346) compared to frequency of an equally sized set of control 8mers randomly sampled. *p-*value computed from a two-sided Mann-Whitney *U* test. (**B**) Positional frequency over 1000 nt, 12-nt bins, of the frequency of aSD-like 8mers in *B. subtilis* CDSs of simple monocistronic TUs. Grey and red lines correspond to the same sets of sequences compared in **A**.

**Figure S9.**
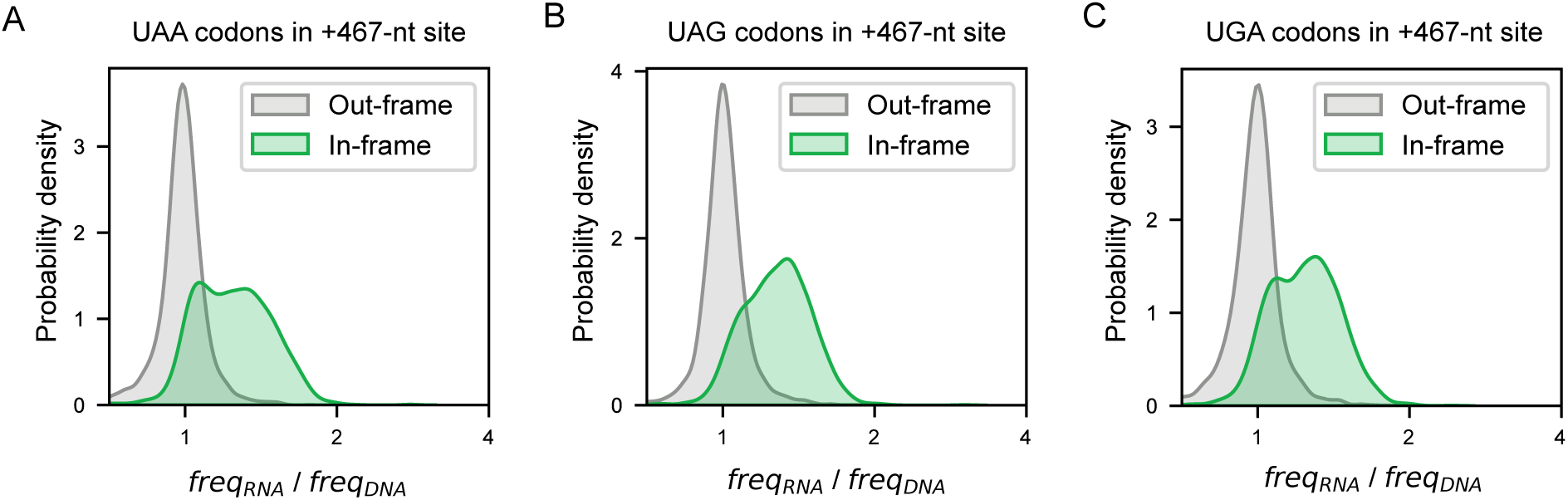
Stabilizing effect of in-frame stop codons at the +467-nt site. (**A-C**) The in-frame and out-of-frame effect on mRNA level of instances of the stop codons UAA (n = 2,015 in, n = 3,981 out) (**A**), UAG (n = 2,022 in, n = 3,960 out) (**B**), and UGA (n = 2,022 in, n = 4,018 out) (**C**) in +467-nt library. SUPPLEMENTAL FILES

## SUPPLEMENTAL FILES

1. Extended Methods

2. Supplementary Table S1- Strains

3. Supplementary Table S2- Plasmids

4. Supplementary Table S3- Oligos

5. Supplementary Table S4- Reagents

