## Extended Methods for "Long-range mRNA folding shapes expression and sequence of bacterial genes"

### GFP REPORTER

#### Reporter/strain description

The GFP reporter (for Figures 5A-F) consists of a Superfolder GFP open reading frame that is under the control of a Ptet promoter that is induced by aTc. This reporter is integrated into the *B. subtilis lacA* locus. The degenerate barcodes (10 bases) are in the 3′ UTR of GFP. The 10mer sequences were computationally collapsed into 8mer sequences for analyses (see 'GFP reporter data analysis').

Figure 5D shows a schematic where the barcode is in a hairpin – from now on referred to as "Barcode 1" – 858 bp downstream of the transcription start site of GFP. A second barcode "Barcode 2" is placed further downstream (891 bp downstream of the transcription start site) and not in a hairpin (which Figure 5A depicts. In Figure 5D, the "Barcode 1" is variable and the barcode in the "Barcode 2" position is a constant 10mer. In Figure 5A, "Barcode 2" is variable and the barcode in "Barcode 1" position is a constant 10mer. The entire 3′ UTR of the GFP reporter is 229 bp long, and "Barcode 1" is in position 36-45 of the 3′ UTR and "Barcode 2" is in position 69 to 78 of the 3′ UTR given base 1 is the base immediately downstream of the stop codon,.

For the variable barcode in either position, a constant sequence barcode (represented by the oligos fixed_barcode_background_1 and fixed_barcode_background_2) was assigned to the other position in the cloning process. The barcodes are first assembled on different vectors (barcode_in_hairpin into barcode_in_hairpin_background_vector and barcode_not_in_hairpin into barcode_not_in_hairpin_background_vector). A sequence containing Superfolder GFP is then inserted into barcode_in_hairpin_background_vector. Subsequently, the vectors for both barcodes are restriction digested separately and purified for the linearized portion containing the barcode. The two linearized barcode vectors are then combined in a cloning step to bring "Barcode 1" and "Barcode 2" into the same plasmid sequence. The complete assembly of the vector is described below.

Furthermore, the strain background is *Bacillus subtilis* 168 with dCas9 integrated into the *amyE* locus. The dCas9 has a pHyperSpank promoter which is inducible by IPTG. A random guide RNA (TGTCAGTACGAAGATACAGG in the seed sequence, which that does not match any sequence in the genome) is also expressed. This strain background was used for a different project, but adapted for use in this paper.

#### Barcode 1 assembly

barcode_in_hairpin_background_vector was linearized using PCR. 32 reactions of 25 µl each were set up using the NEBNext® Ultra™ II Q5® Master Mix (NEB, M0544L), 1 ng of the plasmid, and the primers oJX25_utr_5_backbone_F (0.5 µM) and oJX92_utr_5_backbone_amp_4_R (0.5 µM). The following cycling conditions were used: 98°C for 2 min, 20 cycles (98°C for 10 s, 58°C for 15 s, 72°C for 4 min 30 s), 72°C for 5 min. The 32 PCR reactions were then pooled and the background vector was depleted by addition of 16 µl of Dpn1 (NEB, R0176S) and incubation for 2 hours at 37°C. The products were then purified via a 1X left-side SPRI (Aline Biosciences, PCRClean DX) and eluted in 100 µl water.

barcode_in_hairpin was amplified via PCR using oJX91_utr_5_amp_5_F (0.5 µM) and oJX4_utr_5_amp_R (0.5 µM), and 1 ng of the oligo in twenty-four 25 µl reactions. The following cycling conditions were used: 98°C for 2 min, 12 cycles (98°C for 10 s, 58°C for 15 s, 72°C for 45 s), 72°C for 5 min. The reactions were pooled and cleaned using a 1.8X left-side SPRI and eluted in 100 µl water.

The amplified product was assembled using 4 µg of linearized barcode_in_hairpin_background_vector and 944.9 ng of barcode_in_hairpin (1:10 backbone insertion ratio) using the NEB Hifi DNA Assembly Master Mix (NEB, E2621L) in an 80 µl reaction. The reaction was incubated at 50°C for 1 hour. The assembled product was then purified via a 1X left-side SPRI and eluted in 20 µl water.

The 20 µl elution was split into two 10 µl volumes, each of which was transformed into 100 µl of NEB 10-beta Electrocompetent *E. coli* (NEB, C3020K) via electroporation using 2kV in a 0.1 cm gap cuvette (Bio-Rad, 1652089). 890 µl of recovery media (NEB, B9035S) was added immediately after electroporation. 2 mL additional recovery media was added and the cells were split into 3 separate aliquots, each containing 1 mL, and recovered for 1 hour. Dilution plating estimated a complexity of $7.1 \times 10^8$ transformed cells.

The electroporated library was grown for ~15 hours, spun down at 3220 rcf for 15 minutes at 4°C. Plasmids were extracted from cell pellets using the ZymoPURE II Plasmid Maxiprep Kit (Zymo Research, D4202).

The assembled library was then digested by SfiI (NEB, R0123S). As SfiI requires two copies of recognition sites to cleave, a double-stranded helper oligo (oJX106_SfiI_pgl4_1_73bp_duplex, at 410 ng/µl) was spiked into the reaction. In total, a 200 µl reaction was set up with 5 µl SfiI, 10 µg of the library, and 10 µl of the helper oligo and digested at 50°C for 2 hours. The resulting product was then cleaned using a 1X left-side SPRI and eluted in 50 µl of water.

An amplicon containing GFP was created by PCR and restriction digest. 34 reactions of 25 µl each were set up using the NEBNext® Ultra™ II Q5® Master Mix (NEB, M0544L), 1 ng of original_GFP_plasmid, and the primers oJX34_utr_5_lib_gbl_amp_2_v2_F (0.5 µM) and oJX23_utr_5_lib_gblock_2_R (0.5 µM). The following cycling conditions were used: 98°C for 2 min, 20 cycles (98°C for 10 s, 67°C for 15 s, 72°C for 45 s), 72°C for 5 min. The 34 PCR reactions were then pooled and the background vector was depleted by addition of 16 µl of Dpn1 (NEB, R0176S) and incubation for 2 hours at 37°C. The products were then purified and the background vector further depleted by a 0.4X SPRI and eluted in 100 µl water.

The GFP amplicon was then digested by DraIII (NEB, R3510S) to create sticky ends for ligation and dephosphorylated to prevent self-ligation. 20 µg of the amplicon was incubated with 10 µl DraIII and 38 µl Quick CIP (NEB, M0525S) (1 µl per pmol of DNA) in a 600 µl reaction at 37°C for 1 hour. The reaction was heat inactivated at 80°C for 2 minutes, purified with a 1X left-side SPRI, and eluted in 70 µl water.

The linearized vector was then assembled with the GFP amplicon in a T4 ligase (NEB, M0202T) reaction. 4 µg of the SfiI-digested construct, 1502.7 ng of the GFP construct (1:3 backbone to insert ratio), 4.76 µl of T4 ligase (2,000 units per 200 fmol of vector) in a 70 µl reaction. The reaction was heat inactivated for 10 min at 65°C, cleaned using a 1X left-side SPRI, and eluted in 50 µl of water.

The resulting product was then digested by SfiI again to remove potentially re-ligated background vector backbones. This was accomplished in a 100 µl reaction using the 50 µl elution, 4 µl of the helper oligo, and 2 µl of SfiI. The reaction was incubated at 50°C for 2 hours. The digest was then cleaned using a 1X left-side SPRI and eluted in 50 µl.

Then, a RecBCD (NEB, M0345S) digest was used to remove residual linear vectors. The 50 µl elution from the SfiI digest was mixed with 2 µl RecBCD and 1 mM ATP in a 200 µl total reaction and incubated at 37°C for 1 hour. The reaction was purified using a 1X left-side SPRI and eluted in 20 µl.

The 20 µl elution was split into two 10 µl volumes, each of which was transformed into 100 µl of NEB 10-beta Electrocompetent *E. coli* (NEB, C3020K) via electroporation using 2kV in a 0.1 cm gap cuvette. 890 µl of recovery media was added immediately after electroporation. Per electroporation, 2 mL additional recovery media was added and the cells were split into 3 separate aliquots, each containing 1 mL, and recovered for 1 hour. Each aliquot was expanded in 25 mL of LB with 100 µg/mL carbenicillin. The electroporated library was grown for 8 hours at 225 rpm. The cells were spun down at 3220 rcf for 15 minutes at 4°C. Plasmids were extracted using the ZymoPURE II Plasmid Maxiprep Kit. Dilution plating estimated a complexity of $2.2 \times 10^8$ transformed cells.

The plasmid was then digested with I-SceI (NEB, R0694S) in a 1.2 mL reaction with 80 µl I-SceI, 40 µg of the vector at 37°C for 2 hours. The digest was then cleaned using a 2X left-side SPRI and eluted in 100 µl water. In the elution the SPRI beads were kept for the following step.

The digest yielded two products – a ~1 kb product with GFP and another ~6.5 kb product resulting from the vector. To isolate the GFP vector from the backbone, a specialized SPRI was performed following Stortchevoi et al. 2020. 40 mL of $MgCl_2$ PEG solution was made by mixing 20 mL of 1M $MgCl_2$ (ThermoFisher Scientific, AM9530G), 5 mL 40%, and 15 mL of water. This specialized SPRI was performed by adding 18 µl $MgCl_2$ PEG solution to 100 µl of the solution (containing beads) mixing and incubating for 5-10 minutes, then following the rest of the left-side SPRI protocol normally. This step was repeated 2 times to deplete the vector backbone. Residual small fragments (250-300 bp), likely from vector backbones, were removed by 0.5X left-side SPRI cleanups when present. The final product was eluted in 25 µl water.

For fixed_barcode_background_1, because the oligo is not a library, it was cloned using smaller throughput approach. The oligo was amplified in a 25 µl reaction using the same PCR conditions as barcode_in_hairpin, except with 1 µl of the oligo (1-2 ng). The amplified product was cleaned using a 1.8X left-side SPRI and eluted in 30 µl water. The amplified product was assembled into the backbone using 100 ng of linearized barcode_in_hairpin_background_vector and 24.4 ng of fixed_barcode_background_1 (1:10 backbone to insert ratio) using the NEB Hifi DNA Assembly Master Mix in a 10 µl reaction. The reaction was then used to transform into *E. coli* with the Mix & Go! Competent Cells - DH5 Alpha (Zymo Research, T3009). The transformed cells were plated on plates with carbenicillin and incubated overnight at 37°C to derive colonies. Clones were grown up and sent to Plasmidsaurus for sequence verification. Clones without mutations had plasmids extracted using the ZymoPURE II Plasmid Midiprep Kit (Zymo Research, D4200). The plasmid was then digested by SfiI (100 µl reaction with 5 µg of plasmid, 5 µl of the helper oligo, and 2.5 uL of SfiI at 37°C for 2 hours), 1X left-side SPRI cleaned, and eluted in 35 µl water. A T4 ligase reaction with the SfiI-digest and the GFP digested amplicon then set up reaction and ran under the same conditions, except with half the concentrations and volume as the barcode_in_haripin assembly. The ligase reaction was cleaned with a 1X left-side SPRI and eluted in 25 uL water. A subsequent SfiI digest and RecBCD digest was then done using the same reaction conditions as the barcode_in_haripin assembly. The reaction was cleaned with a 1X left-side SPRI and eluted in 25 uL water. The subsequent product was then diluted and transformed into *E. coli* with the Mix & Go! Competent Cells - DH5 Alpha (Zymo Research, T3009). The transformed cells were plated on plates with carbenicillin and incubated overnight at 37°C to derive colonies. Clones were grown up and sent to Plasmidsaurus for sequence verification. Clones without mutations had plasmids extracted using the ZymoPURE II Plasmid Midiprep Kit.

**Barcode 2 assembly**

barcode_in_hairpin_background_vector was linearized using PCR. 32 reactions of 25 µl each were set up using the NEBNext® Ultra™ II Q5® Master Mix, 1 ng of the plasmid, and the primers oJX8_linearize_utr_5_gblock_F (1 µM) and oJX9_linearize_utr_5_gblock_R (1 µM). The following cycling conditions were used: 98°C for 2 min, 20 cycles (98°C for 10 s, 72°C for 15 s, 72°C for 5 min), 72°C for 5 min. The 32 PCR reactions were then pooled and the background vector was depleted by addition of 16 µl of Dpn1 and incubation for 2 hours at 37°C. The products were then purified via a 1X left-side SPRI and eluted in 100 µl water.

barcode_not_in_hairpin was amplified via PCR using primers oJX6_guide_RNA_amp_3_F (0.5 µM) and oJX7_guide_RNA_amp_R (0.5 µM), and 1 ng of the oligo in 24 25 µl reactions. The following cycling conditions were used: 98°C for 2 min, 12 cycles (98°C for 10 s, 61°C for 15 s, 72°C for 45 s), 72°C for 5 min. The reactions were pooled and cleaned using a 1.8X left-side SPRI and eluted in 100 µl water.

The amplified product was assembled using 4 µg of linearized barcode_not_in_hairpin_background_vector, 1.058 µg of barcode_not_in_hairpin (1:10 backbone insertion ratio) the NEB Hifi DNA Assembly Master Mix in a 120 µl reaction. The reaction was incubated at 50°C for 1 hour. The assembled product was then purified via a 1X left-side SPRI and eluted in 20 µl water.

The 20 µl elution was split into two 10 µl volumes, each of which was transformed into 100 µl of NEB 10-beta Electrocompetent *E. coli* via electroporation using 2kV. 890 µl of recovery media was added and the electroporated cells were recovered for 1 hour. Per electroporation, 2 mL additional recovery media was added and the cells were split into 3 separate aliquots, each containing 1 mL, and recovered for 1 hour. Each aliquot was expanded in 25 mL of LB with 100 µg/mL carbenicillin. The electroporated library was grown for 6 hours at 225 rpm. The cells were spun down at 3220 rcf for 15 minutes at 4°C. Plasmids were extracted using the ZymoPURE II Plasmid Maxiprep Kit. Dilution plating estimated a complexity of $1.47 \times 10^8$ transformed cells.

The assembled library then was digested with SfiI. A 200 µl reaction was set up with 10 µl SfiI, 10 µg of the library, and 10 µl of the helper oligo (at ~410 ng/µl) and digested at 50°C for 2 hours.

To isolate the linearized vector and separate from uncut plasmid and plasmid doublets, a 0.75% agarose gel was cast with 2.5 µg of DNA put into each well (4 wells). The gel was ran for 100V for 2 hours. For each well, a band corresponding to the linearized vector was cut. Each gel piece was split approximately into 1/3$^{rd}$ (12 tubes total) and processed using the Zymoclean Large Fragment DNA Recovery Kit (Zymo Research, D4045) to recover the linearized vector. Each 1/3$^{rd}$ gel piece was incubated with 1 mL buffer ADB from the kit.

For fixed_barcode_background_2, because the oligo is not a library, it was cloned using smaller throughput approach. The oligo was amplified in a 25 µl reaction using the same PCR conditions as barcode_not_in_hairpin, except with 1 µl of the oligo (1-2 ng). The amplified product was cleaned using a 1.8X left-side SPRI and eluted in 30 µl water. The amplified product was assembled into the backbone using 100 ng of linearized barcode_not_in_hairpin_background_vector and 24.4 ng of fixed_barcode_background_2 (1:10 backbone to insert ratio) using the NEB Hifi DNA Assembly Master Mix in a 10 µl reaction. The reaction was then used to transform into *E. coli* with the Mix & Go! Competent Cells - DH5 Alpha (Zymo Research, T3009). The transformed cells were plated on plates with carbenicillin and incubated overnight at 37°C to derive colonies. Clones were grown up and sent to Plasmidsaurus for sequence verification. Clones without mutations had plasmids extracted using the ZymoPURE II Plasmid Midiprep Kit. The assembled vector was then digested with SfiI and the product was subsequently gel extracted in the same manner as barcode_not_in_hairpin, except with only half the digest amount (5 µg, and thus 2 wells containing 2.5 µg were ran in the gel).

**Merging of Barcode 1 vector and Barcode 2 vector**

The following method is used to describe the assembly of both barcode_in_hairpin with fixed_barcode_2 and barcode_not_in_hairpin with fixed_barcode_1. The vector was assembled using the NEBuilder Hifi DNA Assembly Master Mix with 1.5 µg of the SfiI digested fixed_barcode_2/barcode_not_in_hairpin vector and 399 ng of the barcode_in hairpin/fixed_barcode_1 vector containing GFP (2 to 1 vector to insert ratio) in a 50 µl total volume reaction and incubating at 50°C for 1 hour. The resulting digest was then cleaned by 1X left-side SPRI and eluted in 25 µl.

The 25 µl elution was then digest with SfiI to remove potential religated backbone vectors using 1 µl of SfiI and 2 µl of the helper oligo (at ~410 ng/µl) in a 50 µl reaction at 50°C. This digest was then purified by a 1X left-side SPRI and eluted in 50 µl.

Linearized products were removed using a RecBCD digest on the entire elution. 1 µl RecBCD and 1 mM ATP were used in a 100 µl reaction incubated for 1 hour at 37°C The reaction was then cleaned by a 1X left-side SPRI and eluted in 20 µl water.

The 20 µl elution was split into two 10 µl volumes, each of which was transformed into 100 µl of NEB 10-beta electrocompetent cells via electroporation using 2kV. 890 µl of recovery media was added and the electroporated cells were recovered for 1 hour. Per electroporation, 1 mL additional recovery media was added and the cells were split into 2 separate aliquots, each containing 1 mL, and recovered for 1 hour. Each 1 mL was put into 625 mL LB with carbenicillin (100 µg/mL) in a 2.8 L flask and grown for 12 hours. The resulting cells (2.5 L total per plasmid type split across 4 flasks) were then spun down in batches, with each spin occurring at 6000 rpm for 6 min. Plasmids were extracted using the ZymoPURE II Plasmid Gigaprep Kit (Zymo Research, D4204). Dilution plating estimated a complexity of 5.2 x $10^7$ transformed cells for the barcode_in_hairpin plasmid and 3.93 x $10^7$ transformed cells for the barcode_not_in_hairpin plasmid.

**Bacillus transformation of barcode libraries**

200 µg of the library vector was first linearized with 50 µl I-CeuI (NEB, R0699L) in a 2.5 mL reaction for 3 hours at 37°C.

To transform into *B. subtilis*, a colony of dCas9_1 (168 strain with dCas9 integrated using a pDR111_dCas9) was picked and grown in 550 mL LB. The culture was then grown to OD ~1-1.2. Then 1.1 mL of aTc (10 µg/mL, in ethanol) was added and grown for 1 hour 40 minutes. Two 50 mL cultures were spun down at 3220 rcf for 6 minutes at room temperature and the media was discarded and resuspended with 25 mL of pre-warmed 37°C LB. The 25 mL of resuspended culture was aliquoted into 25 mL of LB with ~1.25 mL of the I-CeuI digested DNA in a 1 L flask. Each flask was then grown for 8.5 hours. Afterwards, 10 µl of the culture from each flask was aliquoted for dilution plating and the remaining 50 mL was put in 450 mL LB with 5 µg/mL kanamycin. The cultures were grown for ~12 hours and then both flasks were pooled and frozen in ~20-30 glycerol stocks with 0.625 mL culture and 0.375 mL 40% glycerol in each tube (batch_1).

To further increase library complexity, the transformation process was repeated a second time to freeze down another batch of 20-30 glycerol stocks (batch_2).

**Collection of RNA and DNA from frozen stocks**

One aliquot from batch_1 and one aliquot from batch_2 were thawed and grown in 100 mL LB in a 1L flask for ~3 hours until OD 0.2-0.4. 3 replicates of 560 mL LB culture were prepared with 5 µg/mL Kanamycin and 20 ng/mL aTc in a 2.8 L flask. The OD from each batch was quantified and mixed in a 50/50 proportion based on the OD. Each of the prepared 3 flasks were seeded so that each flask contained 1.56 $x10^5$ cells/mL to ensure 10 doublings into exponential phase. The cells then went through 10 population doublings until an OD of 0.3. Afterwards, for each replicate, two batches of 10 mL of culture were aliquoted into 15 mL tubes on ice for subsequent DNA extraction. 39 mL of culture was aliquoted into 6 mL of Phenol Stop solution (2% Phenol (Sigma-Aldrich, P4557-100ML), 0.01M EDTA, 60% EtOH). All tubes were then spun down at 3220 rcf at 4°C for 5 minutes. The tubes were then flash frozen and stored in -80°C.

Genomic DNA was extracted using the Monarch® Spin gDNA Extraction Kit (NEB, T3010S). Thawed bacteria were then resuspended with 100 µl 50 mM Tris-Cl, 0.5 mM EDTA pH 7.5. 10 µl was of T4 Lysozyme (NEB, P8115) was then added and incubated for 5 minutes at room temperature. Subsequently, 10 µl of Proteinase K (NEB, P8107S) and 100 µl of Tissue Lysis Buffer were added and incubated at 56°C for ~30 minutes in a thermal mixer at 1400 rpm. 3 µl of RNase A was then added, vortexed, and incubated for 5 minutes at 56°C at 1400 rpm. The sample was then subsequently processed using the "Genomic DNA binding and elution steps" from NEB's "Protocol for Genomic DNA Extraction from Gram-positive Bacteria using NEBExpress® T4 Lysozyme (NEB #P8115) and the Monarch Spin gDNA Extraction Kit (NEB #T3010)" (https://www.neb.com/en-us/protocols/2023/09/20/protocol-for-genomic-dna-extraction-from-gram-positive-bacteria-using-nebexpress-t4-lysozyme-neb-p8115-and-the-monarch-genomic-dna-purification-kit-neb-t3010#genomic).

RNA was extracted as follows. The cells thawed and mixed with 300 µl of TE/Lysozyme (Sigma-Aldrich, L6876-10G). 2 mL of RLT/ β-mercaptoethanol (Sigma-Aldrich, M6250-100ML) solution were added and vortexed. 2 mL of 70% ethanol was then added and vortexed. The resulting ~4.3 mL of lysate was then processed using the subsequent manufacturer steps from the Qiagen RNeasy Midi Kit (Qiagen, 75144) (refer to "RNeasy Midi/Maxi Protocol for Isolation of Total RNA from Animal Cells" from the RNeasy® Midi/Maxi Handbook). The sample was eluted in 150 µl and 1 µl of SUPERase•In RNase Inhibitor (ThermoFisher Scientific, AM2694) was added to the sample and frozen at -80ºC.

**Construction of RNA/DNA libraries**

For each replicate, 35 µg of RNA was incubated in DNase digest with 4 µl Turbo DNase (ThermoFisher Scientific, AM2238) in a 100 µl reaction at 37°C for 30 minutes. The reaction was then inactivated by adding 3.09 µl 0.5M EDTA (0.015M final concentration) (ThermoFisher Scientific, AM9260G) and incubated at 75°C for 10 minutes. The samples were then purified via a 1.8X left-side SPRI using RNA SPRI beads (Aline Biosciences, C-1005-500) and eluted in 30 µl. 1 µl of SUPERase•In RNase Inhibitor was added to the elution. Per sample, 20 µg of RNA was then used for Reverse Transcription following Superscript III using random hexamer primers (400 ng) in a 160 µl reaction. The 160 µl was split into two 80 µl reactions and incubated at 50°C for 1 hour. Each 80 µl reaction was then inactivated by adding 16 µl 1M NaOH (diluted from MilliporeSigma, 72068-100ML) and heated at 95°C for 5 minutes. Then,16 µl 1M HCl (Fisher Scientific, SA48500) was added to neutralize the reaction. The resulting sample was then cleaned by a 1.8X left-side SPRI and eluted in 30 µl $H_2O$.

qPCR on the cDNA was then performed with 1.5 µl of sample, primers oJX136_lib_amp_guide_UMI_nextera_R (0.5 µM) and oJX134_MPRA_lib_amp_utr_no_UMI_R (0.5 µM), and 1.7 µl of SYBR Green I (ThermoFisher Scientific, S7563) diluted 1:10000 using the NEBNext® Ultra™ II Q5® Master Mix in a 10 µl reaction to estimate cDNA amount via cycle thresholds. All samples had a cycle threshold between 9-12. 1.5 µl of DNase-digested RNA from all replicates were also ran using the same qPCR setup on the same plate to assess residual plasmids, none were detected (RNA cycle thresholds for all samples were greater than 26).

Two PCRs were then used to build the final libraries for both DNA and cDNA from the reverse transcriptase reactions. The first PCR uses two cycles used to add unique molecular identifiers and sequencing adapters. The second PCR completes the addition of the sequencing adapters and adds sequencing indeces. For the first PCR, for DNA, 4 µg of genomic DNA was used. For the cDNA, 55 µl of the 60 µl elution was used. Each sample was split into 4 separate 100 µl PCR reactions. Each PCR reaction used the NEBNext® Ultra™ II Q5® Master Mix with primers oJX136_lib_amp_guide_UMI_nextera_R (0.5 µM) and oJX134_MPRA_lib_amp_utr_no_UMI_R (0.5 µM). The following cycle conditions were used: 98°C for 1 min, 2 cycles (98°C for 10 s, 66°C for 30 s, 72°C for 1 min), 72°C for 2 min. The PCRs for each sample was then pooled (400 µl total) and cleaned using a 1.8X left-side SPRI and eluted in 100 µl. The sample was then cleaned using a 1.8X left-side SPRI again to remove residual primers from the first PCR and eluted in 165 µl water. ~90% of the elution was then carried over to the 2$^{nd}$ PCR. This PCR also consisted of 4 separate 100 µl PCR reactions with the elution split evenly across the 4 reactions. Each PCR reaction used the NEBNext® Ultra™ II Q5® Master Mix primers Element_index_read_1 (0.5 µM) and Element_index_read_2 (0.5 µM). The following cycle conditions were used: 98°C for 30 s, 2 cycles (98°C for 10 s, 72°C for 30 s, 72°C for 30 s), 72°C for 2 min. The 4 separate PCR reactions were then pooled (400 µl total) and cleaned using a 1.8X left-side SPRI and eluted in 100 µl water. A double-sided SPRI to deplete residual genomic DNA was then performed using 0.5X right-side SPRI to remove large fragments, followed by a 1.8X left-side SPRI and eluted in 35 µl water.

The samples from all replicates were then pooled by concentration to ensure equal representation and sequenced on an Element Aviti system using 2x150 bp sequencing.

**GFP reporter data analysis**

Flash v1.2.11 (Magoč and Salzberg, 2011) was used to merge read 1 and read 2 from sequencing. Perfect matches to expected adapters (17-23 bp) flanking the 10mers were used to locate the 10mer. 10mer read counts were collapsed to 8mers by the following method. For every unique 8mer, the counts for all 10mers with the given 8mer were added together. DESeq2 (Love et al. 2014) was used to obtain the log2FoldChange ratios of RNA over DNA for each 8mer.
